# Fever-range temperature enhances B cells *in vitro* but is dispensable for the germinal center *in vivo*

**DOI:** 10.64898/2026.10.12.757555

**Authors:** Sofiane Mohammed, Zineb Sbihi, Patrick Laignel, Ines Houtia, Sabrina Haddou, Emanuele Procopio, Océane Morales, Grégory Franck, Giusepina Caligiuri, Antonino Nicoletti, Marie Le Borgne

**Affiliations:** Université Paris Cité and Université Sorbonne Paris Nord, INSERM, LVTS, F-75018 Paris, France; Department of Cardiology, Assistance Publique-Hôpitaux de Paris, Hôpitaux Universitaires Paris Nord Val-de-Seine, Site Bichat, Paris, France

**Keywords:** Fever, temperature, humoral immunity, vaccine

## Abstract

Fever is a conserved physiological response to immune activation characterized by elevated body temperature. While hyperthermia enhances innate and T cell immunity, its role in shaping the B cell compartment remains poorly defined, despite B cells’ role in long-term protection. Understanding how temperature modulates B cell physiology is relevant to vaccine strategies, particularly fever management around immunization. Therefore, we investigated the influence of fever-range thermal changes on humoral immunity. To decipher the direct effects, primary B cells from C57BL/6 mice were stimulated *in vitro* under mild hyperthermic conditions (∼1.5 °C increase). Fever-range temperature enhanced B cell activation, facilitated entry into cell division, and increased antigen presentation to CD4+ T cells, while transcriptomic analysis revealed enrichment of pathways involved in protein synthesis and mitochondrial respiration. To determine whether these intrinsic effects translated into altered humoral immunity *in vivo*, we developed a novel tunable implant-based system to induce localized, 48-hour hyperthermia around a draining lymph node in awake mice without altering core body temperature. Following subcutaneous NP-OVA/alum immunization, localized hyperthermia localized hyperthermia did not enhance antigen-specific germinal center B cell expansion, antibody titers or IgG affinity on day 9. Moreover, heating during priming did not affect the recall humoral response after a booster on day 21. Thus, fever-range temperature directly modulates murine B cell function *in vitro* but does not enhance humoral immunity *in vivo*, indicating that the biological consequences of intrinsic B cell thermosensitivity depend on its integration within the multicellular and inflammatory signals shaping germinal center responses.

## Introduction

In response to infection or sterile inflammation, the host often initiates fever, a complex and energy-costly systemic response characterized by a transient elevation of the core body temperature (typically 1-4°C) (Evans et al., 2015). This response is evolutionarily conserved across vertebrates, occurring *via* different physiological and behavioral mechanisms in endotherms and ectotherms, which strongly suggest an adaptive advantage (Kluger, 1991 ; Rakus et al., 2017). Indeed, fever is generally associated with improved survival and faster pathogen clearance (Jiang et al., 2000), supporting the view that elevated temperatures optimize immune functions.

The impact of fever-range temperatures (FRT) on the immune system is multifaceted. It is well established that moderate hyperthermia enhances the functions of innate immune cells, including dendritic cell maturation, antigen presentation capacity (Ostberg et al., 2003), and NK cell cytotoxicity. FRT also potentiates the adaptive lymphocyte response (Ostberg et al., 2007). It enhances lymphocyte trafficking to secondary lymphoid organs by upregulating adhesion molecules on the vascular endothelium, thereby facilitating extravasation (Chen et al., 2006 ; Fisher et al., 2010). Furthermore, temperature directly impacts T cell function and activation: it boosts CD8+ T cell cytotoxicity and cytokine production (Mace et al., 2011 ; Mace et al., 2012), while mechanistically, mild hyperthermia increases membrane fluidity and promotes the clustering of lipid rafts, which can act as thermal sensors (Torok et al., 2014). This promotes the clustering of the T cell receptor (TCR) at the immunological synapse, thereby enhancing signaling kinetics and lowering the activation threshold (Mace et al., 2011 ; Mace et al., 2012 ; Zynda et al., 2015).

These thermally sensitive mechanisms, membrane dynamics and the formation of signaling platforms are fundamental processes shared across lymphocyte activation, suggesting that B cell functions may likewise be intrinsically temperature-sensitive. B cell receptor (BCR) signaling, initiation, and antigen internalization rely heavily on the dynamic reorganization of membrane microdomains (lipid rafts) (Gupta et DeFranco, 2003 ; Pierce, 2002). Given that membrane fluidity and raft organization are temperature-sensitive (Torok et al., 2014), it is plausible that FRT directly impacts B cell activation. Katkere et al. provided mechanistic support for a temperature sensitivity of BCR function, but over a different range than the one studied here. Comparing peripheral (33 °C) with normal-to-febrile (37-39 °C) temperatures, they showed that temperature selectively affects lipid raft-dependent BCR functions, namely signaling, receptor ubiquitination and the subsequent processing of cognate antigen, while leaving raft-independent, clathrin-mediated bulk internalization essentially unchanged. At 33 °C, the surface expression of peptide-MHC class II complexes derived from BCR-mediated processing was profoundly slowed, whereas processing of fluid-phase antigens was unaffected (Katkere et al., 2010). Their design did not separate 37 °C from 39 °C, and therefore does not address whether a febrile shift above core temperature further accelerates these steps, which is the question addressed here. This enhanced presentation capacity is critical for B cells, as it optimizes their ability to solicit essential T cell help, a prerequisite for robust activation and germinal center entry. Beyond membrane dynamics, the subsequent steps of rapid proliferation and differentiation into antibody-secreting cells are highly dependent on massive metabolic reprogramming to fuel growth and antibody production (Boothby et Rickert, 2017 ; Doughty et al., 2006) and dynamic cytoskeletal reorganization for antigen acquisition, synapse formation, and cell division (Harwood et Batista, 2010 ; Yuseff et al., 2013). As metabolic activity and cytoskeletal dynamics are both known to be temperature sensitive, they represent plausible pathways through which temperature could modulate B cell responses. Indeed, moderate hyperthermia can increase metabolic flux, by promoting mitochondrial translation (O’Sullivan et al., 2021), and influence cytoskeletal organization in lymphocytes (Di et al., 1997).

However, predicting the *in vivo* humoral response based solely on *in vitro* intrinsic cellular sensitivities remains challenging. *In vivo*, the generation of a robust antibody response, particularly the establishment of memory, relies on the highly orchestrated microenvironment of the germinal center (Dinarello, 2004 ; Victora et al., 2010). It requires precise spatial and temporal coordination between B cells, T follicular helper cells, and other stromal networks. Evidence from ectotherms supports the thermal sensitivity of the humoral immune response, as studies have demonstrated that warmer environmental temperatures significantly accelerate and enhance antibody production in fish and amphibians (Bisset, 1947, 1948 ; Cushing, 1942). However, translating these findings to mammals remains challenging due to confounding variables in existing studies. Antipyretic-based thermomodulation is problematic, as these drugs exert temperature-independent immunomodulatory effects, including direct inhibition of antibody production by human B cells *in vitro* (Bancos et al., 2009). Therefore, even though some clinical studies have shown that prophylactic acetaminophen administration significantly blunts vaccine-induced antibody titers (Prymula et al., 2009 ; Wysocki et al., 2017), these effects cannot be unambiguously attributed to temperature changes, given the drugs’ direct immunosuppressive properties. Alternatively, preclinical models have utilized environmental heating to induce hyperthermia. However, artificially raising ambient temperature to induce hyperthermia activates compensatory thermoregulatory responses (Nakamura, 2024 ; S. Y. S. Wang et al., 2019), and is associated with systemic physiological alterations, including metabolic changes in mice (James et al., 2023 ; Wen et al., 2021), which may independently modulate immune responses.

To address the ambiguity surrounding the role of temperature in humoral immunity, we undertook a two-step approach. First, we characterized the intrinsic thermal sensitivity of primary murine B cells *in vitro*, focusing on activation, proliferation, and T-B cooperation. Second, to determine the physiological relevance of these mechanisms without the biases inherent in previous models, we engineered and utilized a novel electromagnetic induction system to achieve precise, localized thermal modulation of a draining lymph node during immunization. This approach uniquely allowed us to isolate the effects of localized thermal stress on the establishment and maintenance of the humoral response, free from systemic physiological stress and pharmacological confounders.

## Materials and Methods

### *In vitro* B cell Cultures and Stimulation

Fluorescent surrogate particulate antigens (SPAg, 400 nm diameter) coated with anti-mouse Igκ light chain and either ovalbumin (SPAg-OVA) or bovine serum albumin (SPAg-BSA) were prepared as previously described (Sicard et al., 2016). SPAg concentration was measured with a microplate reader (GloMax, Promega).

B cells were purified from spleen (after red blood cell lysis with Ammonium–Chloride– Potassium (ACK) buffer) or subcutaneous lymph nodes using CD19 microbeads followed by LS column separation (Miltenyi Biotec) or Dynabeads™ Mouse CD43 (Untouched™ B Cells) kit (Invitrogen). Purity was checked by flow cytometry. Purified B cells were cultured in complete RPMI-10 (RPMI 1640 with glutamax, 10% FCS, 50 µM 2-ME, 100 units/mL penicillin/streptomycin) supplemented with mouse interleukin-4 (25 ng/mL, R&DSystems) and interleukin-5 (100 U/mL, Biolegend) in V-bottom 96-well plate. B cells were stimulated with either SPAg-OVA (100/cells), anti-IgM (various concentration, eBioscience) or anti-CD40 (various concentration, eBioscience) antibodies at 37 °C or 38.5 °C, for 1 h to 24 h. B cells were then either directly used for flow cytometry analysis or transcriptomics, or washed and cultured at 37 °C after the initial temperature-dependent stimulation period.

### Proliferation Assay

For proliferation assay, splenic and lymph node B cells were loaded with 5 µM CellTrace Violet diluted in PBS-5% FCS for 5 minutes at room temperature (RT), before quenching for 5 minutes with PBS-5% FCS followed by extensive washing. Cells were then stimulated at 37 °C or 38.5 °C for 24 h with different stimuli, then washed and placed back at 37 °C. Proliferation was analyzed by flow cytometry after 3 days in live cells. Percentage of B cells that entered division (divided B cells) and mean number of divisions per cells that entered proliferation (proliferation index) were calculated using FlowJo.

### Antigen-Presentation Assay

OVA-specific CD4 T cells were purified from spleens of OT-II mice following red blood cell lysis with ACK buffer, by using the Dynabeads™ Untouched™ Mouse CD4 Cells Kit (Invitrogen), then loaded with CellTrace Violet as described for the proliferation assay. Purified lymph node B cells were stimulated for 4 h at 37 °C or 38.5 °C with anti-CD40 and either SPAg-OVA or SPAg-BSA as control, then washed and plated in U-bottom microplates with an equal number of OVA-specific CD4 T cells. Proliferation was analyzed by flow cytometry after 3 days in live CD4 OT-II cells. Percentage of OT-II cells that entered division (divided T cells) and mean number of divisions per cell that entered proliferation (proliferation index) were calculated using FlowJo.

### Transcriptomic Analysis (RNA-Seq)

Purified B cells from lymph nodes of five C57BL/6N mice using CD19 Microbeads (Miltenyi Biotec) were stimulated with SPAg-OVA at 37 °C or 38.5 °C or left untreated at 37 °C. After 4 h, cells were pelleted, washed, and either analyzed by flow cytometry to check activation, or frozen as dry pellets (-20 °C) for RNA extraction. Total RNA was extracted using the RNeasy Micro Kit with on-column DNase digestion (Qiagen). The experiment was independently repeated three times to generate biological replicates. RNA quality was assessed using an Agilent 2100 Bioanalyzer. mRNA-seq libraries were prepared using the TruSeq Stranded mRNA kit (Illumina), and sequencing was performed on an Illumina NextSeq 500 platform at the GENOM’IC platform (Institut Cochin).

Reads were aligned to the mouse reference genome (Mus musculus GRCm38, Ensembl release 101). A filter was applied to exclude lowly-expressed genes (count>10 in at least 3 samples for each gene), and normalization was performed using DESeq2 (R). One sample was excluded based on quality control metrics. The normalized count matrix was imported into Partek Flow software for further analysis. Principal component analysis revealed a batch effect, which was corrected using a general linear model. Gene set enrichment analysis (GSEA) was performed using Reactome and Gene Ontology (GO) gene sets. Gene sets were filtered based on a false discovery rate (FDR) ≤ 0.1 and ranked by normalized enrichment score (NES). Differential gene expression analysis was performed using ANOVA.

### Animals and Housing Conditions

Female and male C57BL/6 mice aged 8-10 weeks were used. OT-II Mouse on a C57BL/6 background were used for T cell purification.

Animals were housed in the LVTS animal facility under controlled environmental conditions: 20 ± 2 °C ambient temperature, controlled relative humidity, and 12 h/12 h light/dark cycle (lights on at 7:00 AM). Standard laboratory chow diet and water were provided *ad libitum*. At least a one-week acclimatization period was implemented before experimental procedures. All investigations on mice conformed to the Directive 2010/63/EU of the European Parliament, and review and approval of the study was obtained from the Comité d’Ethique Paris Nord #121 (APAFIS #9687, #18976, #34536, #44582 and #58886). For *in vivo* local heating experiments, mice were transferred to a climatic chamber maintained at 22 °C five days before immunization and kept there until the end of the heating period.

### Immunization Protocols

In some experiments, mice were anesthetized with isoflurane (2-3% of oxygen) and sterile RFID temperature microchips (UID) were implanted intraperitoneally for wireless animal identification and non-invasive temperature measurement using UID mouse matrix devices at least one week before immunization.

Mice under isoflurane anaesthesia were immunized *via* a subcutaneous injection in the hock of a mix of 50 μg of 4-hydroxy-3-nitrophenylacetyl (NP)-ovalbumin (LGC BioSearch) and either 1 μg (∼40-50 μg/kg) of LPS from *Escherichia coli* O111:B4 (Sigma-Aldrich) or 10 μL of an aluminum hydroxide gel (Alhydrogel adjuvant 2%, InvivoGen), diluted in PBS, for apyretic immunization. Control mice received an injection of PBS. In a separate set of experiments aiming at inducing fever in mice, mice received a subcutaneous hock injection of 50 µg LPS (∼2 mg/kg) from *Escherichia coli* O111:B4 together with NP-OVA.

### Local Heating System

The heating system comprised a water-cooled LC resonant circuit (64 kHz) with external magnetic coil and subcutaneous metallic implants. The system utilized a copper coil (21 cm diameter, 3 turns) with Zero Voltage Switching (ZVS) circuit and a variable voltage supply (0-48 V) for magnetic field modulation. Water cooling prevented environmental heating. Custom 3D-printed cages (Clear V4 resin, Formlabs) minimized magnetic interference.

### Local heating in mice

For local hyperthermia experiments, a novel approach was developed using electromagnetic heating of metallic implants positioned adjacent to target lymphoid tissues. Unless otherwise stated, implants consisted of zinc-plated steel flat ring-shaped objects (thickness 0.7 mm, inside diameter: 4 mm, outside diameter: 8.64 mm, Standers).

Implantation surgery was performed under isoflurane anaesthesia (2-3% in oxygen) with comprehensive perioperative analgesia consisting of buprenorphine administration (0.05 mg/kg, subcutaneous) pre-operatively and 6 hours post-operatively. Following abdominal shaving and sterile preparation, a 2-cm longitudinal skin incision exposed the left inguinal lymph node and surrounding adipose tissue. The metallic object was positioned around one inguinal lymph node and secured to the skin using two 8.0 Prolene sutures. Skin closure was achieved with 6.0 Prolene sutures. A four- to six-week recovery period (26 to 42 days) preceded immunization. In experiments conducted to assess the heating efficiency by infrared thermometry in non-anesthetized mice, the metallic implant was placed under the dorsal skin of the mouse instead.

For implant heating, mice were housed in metal-free cages with food and water *ad libitum*. After a 2-day habituation, radiofrequency exposure (64 kHz) was applied at various voltages (0V: control, 20V, 25V, 30V) for up to 48 hours with continuous cage temperature monitoring. Voltage-dependent thermal output was determined using infrared thermal imaging (FLIR A300) on awake mice, comparing temperatures between implanted and non-implanted zones. Manual rectal temperature measurements served as validation controls when needed.

### Sample Collection and Processing

Animals were anesthetized with intraperitoneal ketamine and xylazine at euthanasia dose (150 mg/kg and 30 mg/kg, respectively). Blood samples were collected into heparinized tubes, either by retroorbital puncture for hematological analysis and flow cytometry, or by intracardiac puncture, followed by sequential centrifugation (1,500 × g for 5 min, then 12,000 × g for 15 min) to isolate plasma, which was stored at −20 °C.

Spleens, bone marrow, and inguinal lymph nodes were mechanically dissociated in cold PBS and filtered through a 100-µm strainer. Red blood cells from spleen and bone marrow were lysed using ACK buffer (Ammonium–Chloride–Potassium).

Inguinal fat pads were finely minced into 1–2 mm fragments in IMDM (Gibco) containing collagenase D (0.1 U/mL; Roche) and DNase I (20 U/mL, Roche). The tissue suspensions were then transferred into C-tubes (Miltenyi Biotec) and processed on a GentleMACS Octo Dissociator with Heaters using the program *37C_cus_m_skin_2*. Following mechanical and enzymatic dissociation, the resulting cell suspensions were filtered through a 70 µm strainer and rinsed with ice-cold IMDM supplemented with 2% FBS to inactivate enzymatic activity. Cells were pelleted by centrifugation and resuspended in PBS for antibody staining.

### Flow Cytometry Analysis

Cell suspensions in 96-well plates were stained with LIVE/DEAD™ Fixable Near-IR (Thermo Fisher, 30 min, 4 °C). Non-specific binding blocked with Mouse FcBlock in Brilliant Stain Buffer (BD Biosciences, 10 min, 4 °C), followed by staining with antibody cocktails (see Tables 1-2) diluted in Brilliant Stain Buffer (20 min, 4 °C). Count Beads (Biolegend) were added for absolute cell quantification. Samples were acquired on a LSRFortessa X-20 cytometer (BD Biosciences) with 4 lasers (355 nm, 405 nm, 488 nm and 633 nm) using BD FACSDiva software. Data analysis was performed using FlowJo v10.

### ELISA

ELISA Serum antibody titers were determined using high-binding 96-well plates (Greiner, High Binding). Plates were coated overnight at 4 °C with NP14-BSA or NP4-BSA (LGC Biosearch, 5 μg/mL) in carbonate buffer (15 mM Na₂CO₃, 35 mM NaHCO₃, pH 9.6). After washing with PBS-0.05% Tween 20, plates were blocked for 2 h at room temperature (RT) with PBS-1% BSA. Serially diluted sera were added for 1 h at RT. HRP-conjugated detection antibodies specific for mouse IgM (Invitrogen) and IgG (SouthernBiotech) were then applied and incubated for 1 h at RT. Plates were developed with TMB chromogen (Thermo Scientific), and the reaction was stopped with TMB stop solution (Thermo Scientific). Optical density was measured at 450 nm using a microplate reader (GloMax, Promega). Antibody titers were calculated using the mean value of negative controls plus two standard deviations as the detection threshold. The affinity index was calculated as the ratio of the anti-NP4 to the anti-NP14 IgG titer measured on the same serum: NP4-BSA is bound only by high-affinity antibodies, whereas NP14-BSA is bound by the whole anti-NP repertoire, so that a higher index reflects a larger proportion of high-affinity clones (Brownlie et al., 2008).

### ELISPOT Assay

MultiScreen-IP plates (Millipore) were ethanol-wetted, rinsed with water and PBS, and kept moist throughout processing. Wells were coated with NP14–BSA (LGC Biosearch, 10 μg/mL) in a carbonate buffer (15 mM Na₂CO₃, 35 mM NaHCO₃, pH 9.6) and incubated overnight at 4 °C. Plates were then washed with PBS and PBS-0.05% Tween 20 (PBST) and blocked for 2 h with PBS-BSA 1% at room temperature. Cells were counted, pelleted in V-bottom plates, and resuspended in R15 medium (RPMI 1640 + Glutamax with 15% FBS, 50 μM β-mercaptoethanol, 20 mM HEPES pH 7.0, 1 mM sodium pyruvate, 100 units/mL penicillin/streptomycin) to the appropriate concentration before performing serial fivefold dilutions. Diluted cells were then transferred to the coated ELISPOT plates, which were incubated overnight at 37 °C. For detection, plates were washed with PBS and PBS-T, incubated with HRP-conjugated donkey anti-mouse IgG antibodies (20 ng/mL in PBST-BSA 1%, Invitrogen), washed again, and developed with freshly prepared AEC substrate (BD Biosciences). The reaction was stopped by rinsing with tap water. Plates were air-dried in the dark and stored until analysis. Spot enumeration was performed manually after digital microscope imaging (VHX-X1, Keyence) using FIJI ImageJ.

### Experimental Design and Statistical Analysis

All statistical analyses were performed in JMP Pro 19.0 (SAS Institute Inc., Cary, NC, USA), independently of the graphical visualizations, which were prepared in GraphPad Prism 11 (Dotmatics, Boston, MA, USA).

#### In vitro experiments

For each independent *in vitro* experiment, B lymphocytes isolated from a pool of mice were divided and simultaneously incubated at 37 °C and 38.5 °C, with the paired 37 °C condition serving as the reference condition for that experiment. When multiple technical replicates were performed (1 to 4 per condition), their raw values were averaged prior to any further calculation. For each independent experiment, the response to fever-range temperature was expressed as the ratio of the value measured at 38.5 °C to that measured at 37 °C (38.5 °C/37 °C). The ratio was then log_2_-transformed, and a one-sample t-test was used to test whether the mean log2-ratio of the independent experiments differed from zero (equivalent to a ratio of 1). Results are reported as the fold changes to the paired 37 °C condition. Because the proliferation index is bounded at a minimum of 1 by definition, ratios for this variable were calculated on (proliferation index − 1), representing the average number of divisions beyond the first division, before log2-transformation. Statistical analyses were performed using JMP Pro 19.0. A p-value <0.05 was considered significant.

#### In vivo Experiments

All analyses used linear mixed-effects models to account for the two-batch blocked design of each *in vivo* study. Lognormal variables (normalized alive, B, GC and NP⁺ counts in draining lymph nodes and spleen; IgG and IgM ELISA titers) were log₁₀-transformed before modeling. Mice with undetectable NP⁺ responses were treated as missing and shown as non-responders on the raw-data figures. For the primary vaccination (Figure 3; n = 40, 10 per group), each variable was modeled as Y ∼ group + (1|batch), the four treatment groups (PBS, PBS+Heat, Vaccine, Vaccine+Heat) being the fixed effect. As there was no difference between the PBS and PBS+Heat group, only the PBS group is shown on Figure 3. For the memory experiment (Figure 4; n = 38), single-time point FACS variables were modeled as Y ∼ sex × heat + (1|batch), and repeated serology (IgG, IgM at D21 and D28) as Y ∼ sex × heat × timepoint + (1|mouse) + (1|batch), IgG and IgM in separate models. The log₁₀ %NP⁺ GC variable (Figure 4), which did not converge, was analyzed by two-way ANOVA (sex × heat). Models were fitted by REML with Kenward-Roger degrees of freedom; pairwise comparisons used the Tukey-Kramer adjustment, performed on the highest-order significant interaction. Variance components are reported as a percentage of total variance with Wald 95% confidence intervals. Significance was set at α = 0.05; 0.05 <p <0.10 is reported as a trend, and adjusted p-values below 10⁻⁴ as p <0.0001.

**Figure 1.**
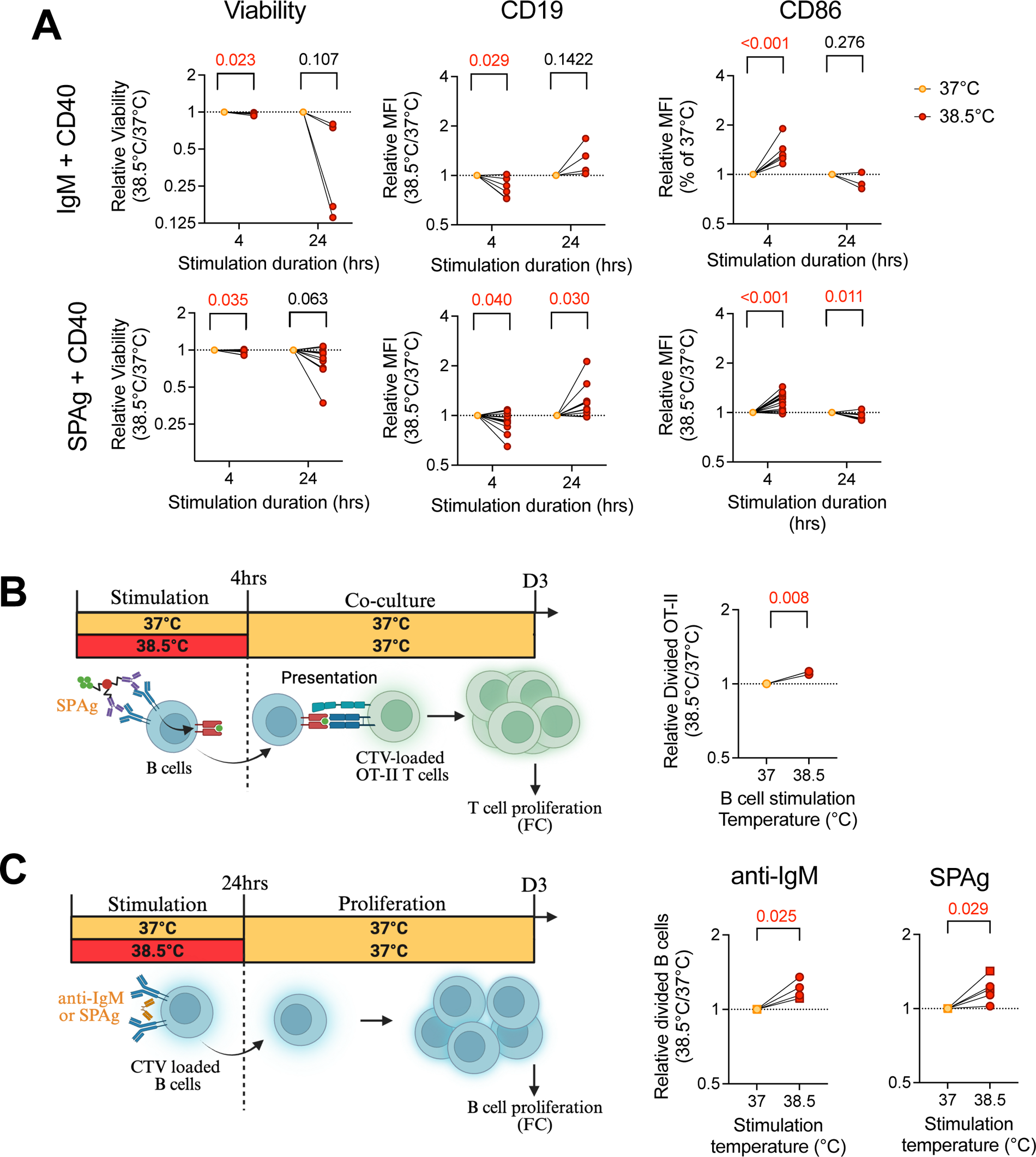
Fever-range temperatures enhance B cell activation and proliferation *in vitro*. (A) Viability and CD19 and CD86 surface expression of B cells stimulated for 4 hrs or 24hrs at 37 °C or 38.5 °C with anti-IgM + anti-CD40 antibodies (top) or SPAg-OVA + anti-CD40 antibody (bottom), assessed by flow cytometry. MFI: median fluorescence intensity, normalized to 37 °C. (B) Experimental procedure for antigen presentation assay (left), and percentage normalized to 37 °C of divided OVA-specific OT-II T cells in coculture with B cells previously stimulated at different temperatures by SPAg-OVA + anti-CD40 (right). (C) Experimental procedure for B cell proliferation assay (left), and percentage of divided B cells (normalized to 37 °C) after initial stimulation at 37 °C or 38.5 °C. In (A-C), each dot represents one independent experiment. Numbers indicate the p-values from one-sample t-tests performed on log2-transformed within-experiment ratios. P-values ≤ 0.05 are shown in red. FC: Flow cytometry. Experimental schematics were created in BioRender. Le borgne, M. (2026) https://BioRender.com/fl9xo4x.

**Figure 2.**
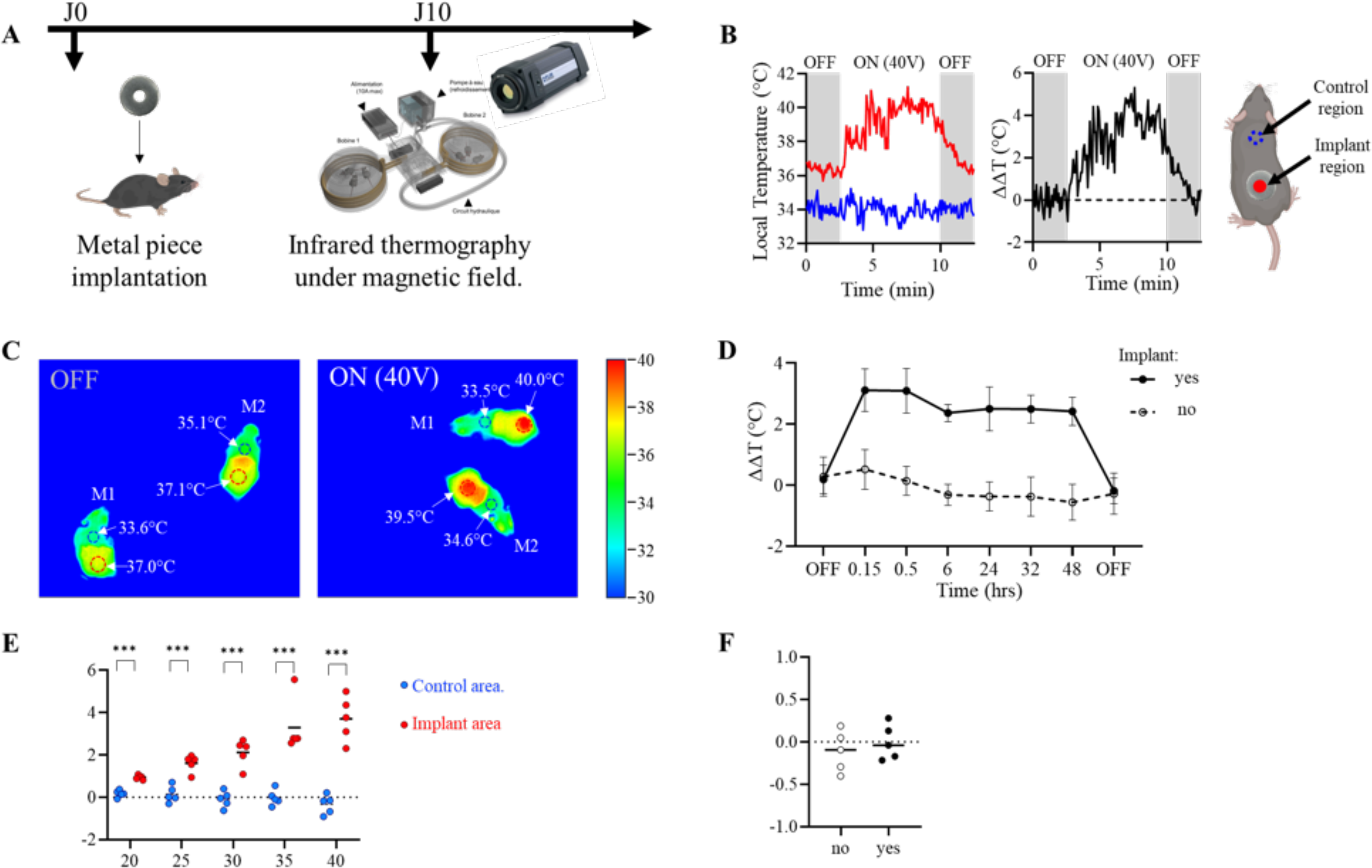
Development and validation of a tunable, localized electromagnetic heating system for *in vivo* lymph node hyperthermia. (A) Schematic of the heating system. A metallic ring-shape object implanted subcutaneously is heated by a radiofrequency electromagnetic field (64 kHz) generated by an external copper coil. Skin surface temperature is monitored by infrared thermography. (B) Representative heating and cooling kinetics at the implant area upon field activation and deactivation (40 V) in awake mice. Absolute skin temperature (left) and corrected temperature ΔΔT = (implant area – control area)_Tx – (implant area – control area)_T0 (right) are shown. (C) Representative infrared thermal images from **movie 2** of two implanted mice (M1, M2) with the electromagnetic field OFF or ON (40 V). Dashed red and blue circles indicate the implant area and the interscapular control area, respectively. Color-coded scale bar from 30 to 40 °C. (D) Thermal stability of localized heating during 48 h of continuous electromagnetic exposure (25 V). Each line represents one mouse (mean ± SD of replicate measurements per time point). Data expressed as ΔΔT as defined in (B). Representative of 2 independent experiments. (E) Differential temperature elevation (ΔΔT) at the implant *versus* control area as a function of applied voltage, measured by infrared thermometry in awake mice. Temperature measurements were analyzed using a linear mixed-effects model with body zone, voltage, and their interaction as fixed effects, and mouse identity and recording (nested within mouse) as random effects, revealing significant effects of body zone (p = 0.0003), voltage (p <0.0001), and their interaction (p <0.0001). Differences in temperature between body zones at each voltage were assessed by pairwise comparisons of least-squares means. (F) Core body temperature in implanted (n=5) versus non-implanted (n=5) mice during electromagnetic field cycling (25V ON/OFF, assessed hourly over 24 h). Each point represents the mean ΔT (mean ON – mean OFF) per mouse across 3 independent measurement sessions. Data were analyzed using a linear mixed-effects model with implant status as a fixed effect, and mouse identity and measurement session as crossed random effects (p = 0.473).

**Figure 3.**
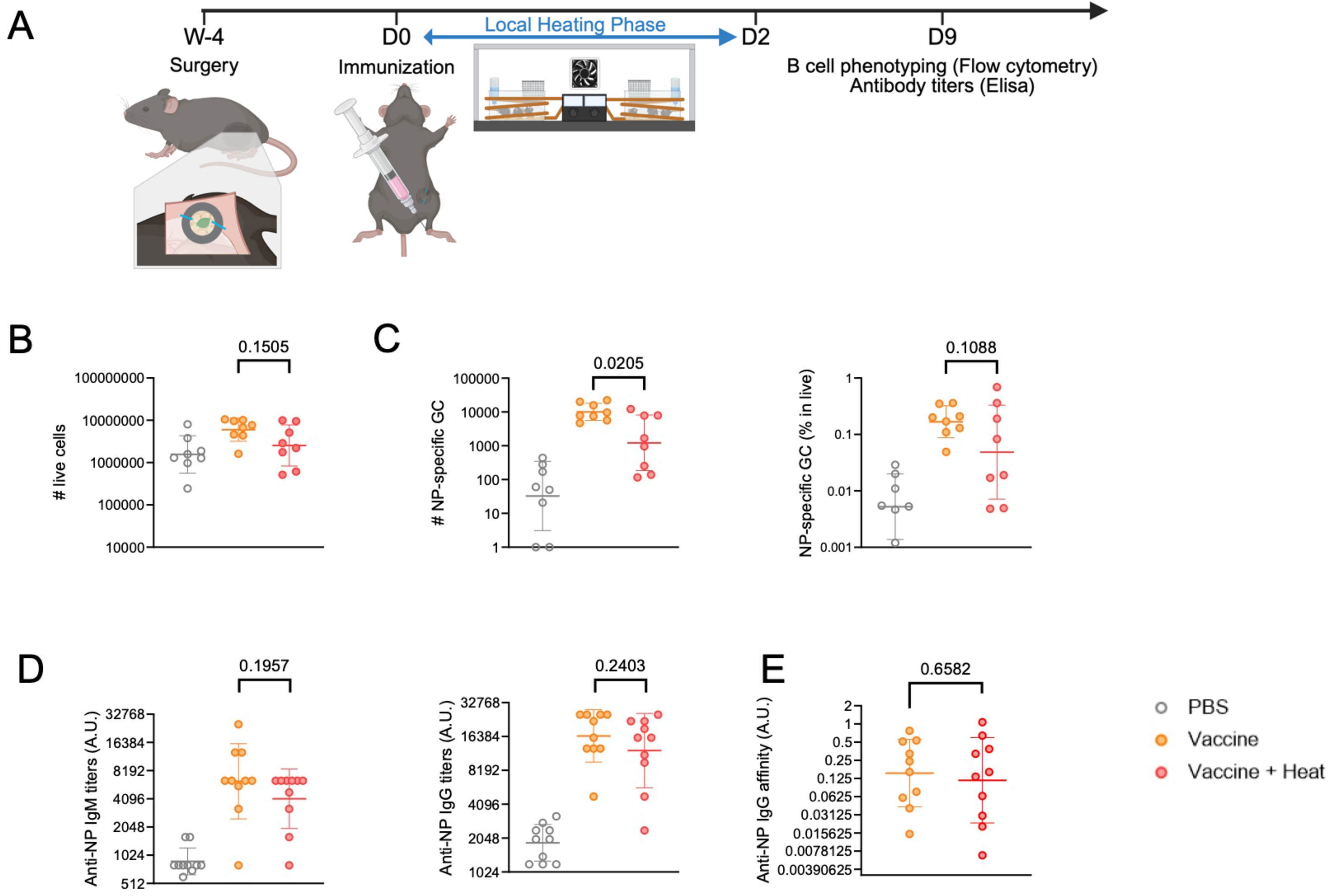
Localized hyperthermia does not augment the primary humoral response to immunization. (A) Experimental design. C57BL/6 mice were implanted with a metallic ring around the left inguinal lymph node 4 to 6 weeks before immunization. At D0, mice received a subcutaneous hock injection of NP-OVA + alum or PBS. Localized electromagnetic heating (+1.5 °C at the implant area, 25 V) was applied from D0 to D2. Draining inguinal lymph nodes and serum were collected at D9. Created in BioRender. Le borgne, M. (2026) https://BioRender.com/fl9xo4x. (B-C) Absolute number of total live cells (B) and absolute number (left) and frequency among live cells (right) of NP-specific germinal center (GC) B cells (C) in the draining inguinal lymph node at D9. Gating strategy as in **Supplementary** Figure 5A. (D-E) Serum NP-specific IgM (left) and IgG (right) antibody titers (D), and IgG affinity index (E), at day 9 post-immunization. Values are plotted on a log2 scale. PBS (grey), Vaccine (orange), Vaccine + Heat (red). Each point represents one mouse. Bars indicate geometric mean ± geometric SD. Data pooled from 2 independent experiments. Statistical comparisons: Tukey-Kramer test. ns, not significant.

**Figure 4.**
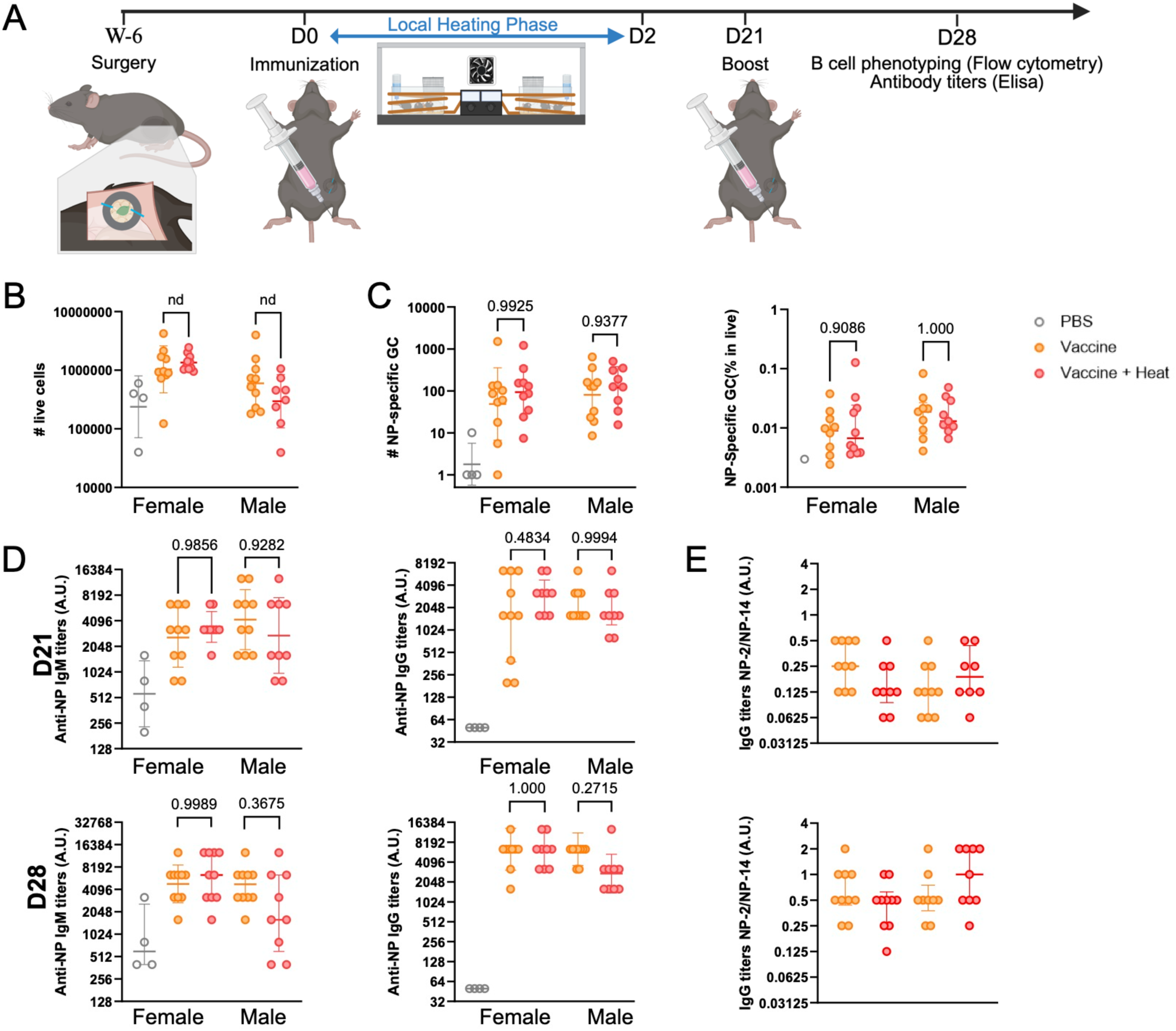
Transient localized hyperthermia during priming does not impact the recall humoral response. (A) Experimental design. Implanted C57BL/6 mice were primed at D0 with NP-OVA + alum or PBS (s.c. in the hock) and received localized electromagnetic heating (+1.5 °C, 25 V) from D0 to D2, as in Figure 3A. A booster immunization with NP-OVA alone (no adjuvant) was administered at D21 in the same hock. Mice were analyzed at D28. Created in BioRender. Le borgne, M. (2026) https://BioRender.com/fl9xo4x. (B-C) Absolute number of total live cells (B) and absolute number (left) and frequency among live cells (right) of NP-specific germinal center (GC) B cells (C) in the draining inguinal lymph node at D28. Gating strategy as in **Supplementary** Figure 5A. (D) Serum NP-specific antibody response. Left: anti-NP14 IgM titers at D21 (top, pre-boost) and D28 (bottom, post-boost). Right: anti-NP14 IgG titers at the same time points. (E) IgG affinity index at D21 (top) and D28 (bottom). Pooling both sexes and both groups, the index increased 4-fold between D21 and D28 (n = 38 paired mice, Wilcoxon signed-rank test, p = 1.6 × 10^−6^), whereas it did not differ between heated and non-heated mice at either time point (Mann-Whitney test, p = 0.41 at D21 and p = 0.56 at D28). PBS (grey), Vaccine (orange), Vaccine + Heat (red). Males (filled circles) and females (open circles) are represented separately within each group. Each point represents one mouse. Bars indicate mean. Data pooled from 2 independent experiments. Statistical comparisons: linear mixed-effects models (fixed effects: heat, sex; random intercept: batch; for the D21 and D28 IgM and IgG serology, timepoint was added as a fixed effect and mouse as a random intercept), with Tukey-Kramer post-hoc; the non-converging log₁₀ % NP⁺ GC variable was analyzed by two-way ANOVA (sex × heat). ns, not significant.

## Results

### Moderate Hyperthermia Enhances Primary B cell Activation *In vitro*

We first sought to determine the direct effects of fever-range temperatures (FRT) on primary B lymphocytes. Splenic B cells were isolated and stimulated *in vitro* under normothermic (37 °C) or moderate hyperthermic (38.5 °C) conditions using either soluble anti-IgM or synthetic particulate antigen (SPAg, (Sicard et al., 2016)) to engage the BCR, both in the presence of an agonistic anti-CD40 antibody as a co-stimulatory signal. FRT reduced cell viability in a time-dependent manner, with a modest effect at 4 hours that became more pronounced at 24 hours relative to 37 °C **(Figure 1A)**. In contrast, it enhanced early B cell activation. Relative to 37 °C, hyperthermia induced a stronger downregulation of CD19 and upregulation of CD86 after 4 hours **(Figure 1A)**, consistent with accelerated activation kinetics observed within the first 0– 6 hours of stimulation **(Supplementary Figure 1A)**. These phenotypic changes suggest that FRT could enhance antigen presentation to T cells and may also promote B cell proliferation. To assess the impact of FRT on antigen presentation, primary B cells were stimulated at 37 °C or 38.5 °C for 4 hours using ovalbumin (OVA)-coated SPAg (SPAg-OVA) , then co-cultured at 37 °C with OVA-specific OT-II CD4⁺ T cells, such that only B cells were exposed to FRT. B cells pre-activated at 38.5 °C induced greater OT-II T cell proliferation at day 3 compared to 37 °C controls **(Figure 1B and Supplementary Figure 1B)**. These results suggest that FRT enhances the ability of B cells to present antigen to CD4⁺ T cells.

To evaluate whether FRT also impacts B cell proliferation, we quantified B cell proliferation following 24 hours of stimulation at 37 °C or 38.5 °C. Hyperthermic conditions increased the fraction of cells entering division (by ∼20% relative to 37 °C) **(Figure 1C)**, whereas the proliferation index, which reflects the average number of divisions undergone by responding cells, was only marginally affected at day 3 **(Supplementary Figure 1C)**. These results indicate that FRT facilitates B cell entry in cell division.

To explore the molecular mechanisms driving the effects of FRT on B cell function, we performed transcriptomic analysis (RNA-Seq) on B cells cultured for 4 hrs at 37 °C versus 38.5 °C. Differential expression analysis revealed differential expression in 257 genes **(Supplementary Figure 1D)**. Gene Set Enrichment Analysis (GSEA) revealed marked changes in pathways related to cellular metabolism and protein synthesis. Hyperthermia induced a significant enrichment of gene sets associated with translational processes, including mRNA capping, mRNA stability (via AU-rich element binding), and ribosomes **(Supplementary Figure 1E)**. In parallel, pathways linked to mitochondrial function were enriched, notably those involved in respiratory electron transport, suggesting enhanced energetic metabolism under FRT conditions **(Supplementary Figure 1E)**.

Altogether, these *in vitro* data indicate that moderate hyperthermia is associated with transcriptional programs supporting increased protein synthesis and mitochondrial metabolism. These changes may contribute to the enhanced proliferation and antigen-presenting capacity observed under FRT conditions.

### Development and Validation of a Tunable Localized Heating System

Considering the direct effects of hyperthermia on B cells *in vitro*, we hypothesized that physiological fever could directly boost the humoral response to T-dependent antigens *in vivo*. Standard pyrogenic stimuli, such as LPS, are known to induce dose- and ambient-temperature-dependent thermoregulatory responses in mice (Romanovsky et al., 2005 ; Rudaya et al., 2005). In our hands, LPS injected subcutaneously at is 2.0 to 2.5 mg/kg together with NP-OVA induced a biphasic rise in core body temperature over the first 48 hours **(Supplementary Figure 2)**. The two waves, between 3 and 7 hours and between 22 and 32 hours, were separated by a return to baseline and reached a 1.4 °C increase in temperature. Therefore, we engineered a novel electromagnetic induction system designed to safely and precisely generate a fever-range hyperthermia for an extended period of time in non-anesthetized animals, strictly localized to a draining lymph node, thereby limiting heat exposure to the targeted tissue and preventing indirect systemic effects that could secondarily influence lymph node responses. This system couples an external radiofrequency magnetic field generator (64 kHz) with a surgically implantable ring-shaped metallic structure that can be placed around the inguinal lymph node **(Figure 2A)**.

We first characterized the heating dynamics of the metallic objects *ex vivo*. Upon radiofrequency exposure, the object temperature increased rapidly, reaching a stable plateau within approximately 5 minutes, and dissipated promptly upon field deactivation **(Supplementary Figure 3A)**. The induced heating was remarkably stable during prolonged continuous exposure (48 hours) without detectable thermal drift **(Supplementary Figure 3B)**. Furthermore, the thermal output was tunable, correlating directly with both the object size and the applied magnetic field strength (voltage) **(Supplementary Figure 3A and 3 C and movie 1),** and reproducible, with low variability between recordings (SD = 0.07 °C) and objects (SD = 0.17 °C) **(Supplementary Figure 3D)**. Horizontal position of the objects had a larger effect, increasing from 1.19 at the center to 2.08 °C at the cage periphery **(Supplementary Figure 3E**), while vertical position had a more modest effect, with the temperature increase ranging from 1.14 to 1.62 °C across the cage height **(Supplementary Figure 3F)**.

As we could induce temperature increases within the range of moderate fever, we next evaluated the thermal performance of the system *in vivo*. Ring-shaped metallic objects were surgically implanted subcutaneously in the mouse’s back to allow infrared thermal imaging of awake, freely moving mice. Electromagnetic exposure induced a strictly localized and reversible temperature elevation at the implant site **(Figure 2B-C and movie 2),** which remained stable over a 48-hour period **(Figure 2D)** and was precisely tunable **(Figure 2E)**. Importantly, this localized thermal stress did not result in systemic temperature changes: rectal temperature measurements taken over 24 hours of continuous local heating showed no significant difference between implanted and sham-operated mice (p = 0.473) **(Figure 2F)**.

Furthermore, we evaluated whether the implant itself, in the absence of heating, altered local or systemic immune homeostasis when implanted around an inguinal lymph node. Blood counts and splenic immune populations were comparable between implanted and sham mice from 15 to 32 days after surgery (**Supplementary Figure 4 A-D)**. In the draining inguinal lymph nodes, we observed a transient increase in B cell numbers peaking around day 22–24 post-implantation, which fully resolved by day 32 **(Supplementary Figure 4F)**. A concomitant macrophage infiltration was detected in the perilymphatic adipose tissue, consistent with a transient foreign body reaction **(Supplementary Figure 4E)**. Critically, no alterations were detected in the contralateral lymph nodes, spleen, or blood at any time point, confirming the strict local nature of this transient reaction. We also assessed if the presence of the metallic implant around the lymph node altered the establishment of a local immune response upon antigen exposure. Implanted or sham-operated mice were immunized subcutaneously with NP-OVA + LPS in the hock, in the absence of electromagnetic field activation, and B cell response was analyzed after 9 days. No significant differences in total CD19+ B cells, GC B cells (B220+ CD19+ IgD-CD95+ CD38low), NP-specific GC B cells (B220+ CD19+ IgD-CD95+ CD38low NP+), or plasmablasts (CD138+) were detected between sham and implanted mice in the draining inguinal lymph node, despite a robust induction of the response in both immunized groups compared to the PBS-injected sham controls **(Supplementary Figure 5)**. These data confirm that the metallic implant *per se* does not impair the magnitude of the primary humoral response.

Altogether, these data validate a stable, tunable, and localized *in vivo* hyperthermia platform that enables selective heating of a single lymph node without systemic effects, providing a controlled framework to investigate the impact of fever-range temperatures on humoral immune responses.

### Localized Hyperthermia does not enhance the Primary B cell Response to immunization

To investigate the functional impact of localized hyperthermia on the humoral response *in vivo*, mice were immunized in the hock with a non-pyrogenic adjuvanted vaccine (NP-OVA + alum). Localized heating (+1.5 °C, 25V) was applied to the draining inguinal lymph node for the initial 48 hours post-immunization **(Figure 3A)**. We analyzed the primary response at day 9, when the germinal center (GC) reaction is well established (Victora et Nussenzweig, 2012). In contrast with the enhancement of B cell function observed *in vitro*, localized heating did not increase the numbers of antigen-specific GC B cells (B220+ CD19+ IgD-CD95+ CD38low NP+) **(Figure 3C)**. Mean NP+ GC B cell numbers were numerically lower in heated mice, likely reflecting both a lower overall leukocyte accumulation in draining lymph nodes and a reduced proportion of NP+ GC B cells, although neither trend reached statistical significance individually **(Figure 3B-C)**.

Consistent with this, serum titers of NP-specific IgM and IgG, reflecting the overall magnitude of the primary humoral response, were comparable between groups **(Figure 3D)**, as was the affinity of the NP-specific IgGs, measured as the ratio of anti-NP4 to anti-NP14 titers **(Figure 3E)**. Altogether, these data indicate that a sustained, early, localized thermal stress does not detectably enhance either the magnitude or the quality of the primary humoral response.

### Localized Hyperthermia Does Not Impact the Memory Humoral Response

Although localized hyperthermia did not detectably alter the GC response measured at its peak, the 48-hour heating window overlaps with the early stages of B cell differentiation that contribute to the subsequent establishment of long-term plasma cell and memory B cell compartments (Elsner et Shlomchik, 2020 ; MacLennan et al., 2003 ; Roco et al., 2019 ; Weisel et al., 2016), raising the possibility that an early thermal perturbation could affect immune memory without altering the magnitude of the primary response measured at day 9. To test it, mice that had received the initial NP-OVA + Alum immunization and the 48-hour localized heating at day 0 were given a booster immunization of NP-OVA (without adjuvant) at day 21 **(Figure 4A)**.

We evaluated the recall response 7 days post-boost (day 28). The initial application of localized hyperthermia did not result in any significant differences in the absolute number or frequency of total and NP-specific GC B cells present in the draining lymph node 7 days after the boost, in either male or female mice **(Figure 4B-C)**. Consistently, the frequency of NP-specific IgG-secreting antibody-secreting cells (ASCs) in the bone marrow and spleen at D28, assessed by ELISPOT in a subset of mice, was unchanged by the early thermal stress **(Supplementary Figure 6)**. Similarly, titers of NP-specific IgM and IgG measured just before the boost (day 21) and post-boost (day 28) were comparable across all groups **(Figure 4D).** The affinity of the NP-specific IgGs was likewise unaffected, both before and after the boost **(Figure 4E)**; across all groups this index increased fourfold between D21 and D28, confirming that affinity maturation proceeded normally. Therefore, in this specific vaccination strategy, a transient early localized hyperthermia does not modulate the establishment nor the recall capacity of the memory humoral response.

## Discussion

The physiological role of fever in optimizing the immune response has long been appreciated, and previous work has documented temperature sensitivity across innate immune cells, T lymphocytes, and lymphocyte trafficking (Evans et al., 2015). In contrast, whether fever-range hyperthermia directly shapes the B cell-driven humoral response *in vivo* has remained difficult to determine, as fever is normally inseparable from the complex inflammatory and physiological changes that accompany it. In this study, we dissected the impact of fever-range temperatures on B lymphocytes, combining *in vitro* characterization with a novel *in vivo* model of localized, drug-free heating of a single draining lymph node to isolate the contribution of temperature itself. Our findings indicate that the intrinsic thermosensitivity of B cells observed *in vitro* does not necessarily translate into enhanced humoral immunity *in vivo* under the conditions tested.

Our *in vitro* findings extend previous observations that B cells are intrinsically responsive to fever-range (+1.5 °C) temperature elevation. Beyond the raft-dependent processing described between peripheral and normal-to-febrile temperatures by Katkere et al. (Katkere et al., 2010), we show that fever-range temperatures also promote early B cell activation and facilitate entry into cell division. These functional changes were accompanied by a transcriptional program enriched for pathways involved in protein translation and mitochondrial respiration, consistent with the increased biosynthetic and energetic demands associated with B cell activation (Boothby et Rickert, 2017). Interestingly, the effects observed in B cells parallel those previously described in other immune populations. Fever-range temperatures have been shown to promote dendritic cell maturation, migration and antigen-presenting function (Ostberg et al., 2003), while enhanced mitochondrial translation has been identified as a key mechanism by which elevated temperature supports the metabolic fitness and effector differentiation of CD8⁺ T cells (Mace et al., 2011 ; O’Sullivan et al., 2021). Our findings therefore demonstrate that B cells are among the growing number of immune cell populations intrinsically equipped to sense and respond to fever-range thermal stress.

The absence of a measurable enhancement of the humoral response *in vivo* despite the intrinsic thermosensitivity of B cells highlights the difficulty of extrapolating cell-intrinsic mechanisms to complex immune responses. A similar discrepancy has recently been described for T lymphocytes, where the beneficial effects of fever-range temperatures observed *in vitro* are largely buffered *in vivo* through dendritic cell crosstalk and IL-12 signaling (Umar et al., 2020). Our own *in vitro* data further indicate that fever-range hyperthermia exerts both stimulatory and detrimental effects on B cells, enhancing early activation while reducing cell viability. Together, these observations suggest that the net impact of temperature on humoral immunity results from the integration of multiple, and sometimes opposing, mechanisms operating within a multicellular network, such that the functional output reflects not only the intrinsic properties of B cells but also their interactions with T follicular helper cells, follicular dendritic cells and the stromal microenvironment that collectively regulates germinal center dynamics (Allen et al., 2004 ; Denton et Linterman, 2017 ; Stebegg et al., 2018 ; Victora et al., 2010).

Beyond tissue-level buffering, another important consideration concerns the nature of the thermal stimulus itself. Physiological fever is not solely defined by an elevation in temperature but constitutes a coordinated systemic response in which thermal changes occur together with pyrogenic cytokines, lipid mediators and widespread vascular and metabolic adaptations (Evans et al., 2015 ; Kluger, 1991 ; Rakus et al., 2017). By design, our model isolates the thermal component from these accompanying signals. While this reductionist approach represents a major strength for dissecting the specific contribution of temperature, it also raises the possibility that the beneficial effects of fever on humoral immunity (Prymula et al., 2009 ; Wysocki et al., 2017) emerge from the integration of thermal and inflammatory cues rather than from temperature alone.

Several features of our experimental model may have further limited the detection of a thermal benefit. The NP-OVA/alum immunization model is characterized by a highly immunodominant and relatively restricted B cell response in C57BL/6 mice, in which affinity maturation follows well-defined stereotypic pathways (Allen et al., 1988 ; Cumano et Rajewsky, 1985, 1986). In addition, alum efficiently promotes germinal center formation and humoral immunity through the induction of robust T cell-dependent responses (Pulendran et al., 2021). Together, these characteristics may place the humoral response close to its maximal magnitude, leaving limited scope for a moderate enhancement induced by fever-range temperatures. This interpretation is further supported by the strong immunodominance of NP over the OVA carrier, which restricts the expansion of subdominant B cell clones (Silva et al., 2017). Whether thermal modulation becomes more apparent under conditions of limiting antigen availability, weaker adjuvanticity or greater clonal diversity remains an important question for future studies.

An additional limitation relates to the temporal profile of the thermal stimulus. To isolate the effect of temperature itself, we applied a sustained, localized +1.5 °C elevation during the first 48 h following immunization. Although this approach reproduces the magnitude of a moderate fever while avoiding pharmacological (Bancos et al., 2009) and systemic confounders, physiological fever is a dynamic process, with body temperature continuously adjusted by central thermoregulatory circuits and inflammatory mediators rather than maintained at a fixed level (Kluger, 1991 ; Nakamura, 2024 ; Rakus et al., 2017 ; T. A. Wang et al., 2019). Whether immune cells integrate transient or fluctuating thermal cues differently from sustained exposure remains unknown.

Beyond the present study, the localized heating platform developed here provides a versatile experimental tool to investigate the role of temperature in immune regulation *in vivo*. Although applied here to a draining lymph node, the approach is readily adaptable to other anatomical sites through implantation of the heating element within or adjacent to the tissue of interest. This opens the possibility of investigating how localized temperature modulation influences immune responses in diverse immunological niches, including tumors (Evans et al., 2015) and chronically inflamed tissues, where tertiary lymphoid organs are known to develop in different contexts (Guillaume et al., 2025 ; Le Borgne et al., 2015).

## Conclusion

In conclusion, our study demonstrates that B cells are intrinsically responsive to fever-range hyperthermia, with enhanced activation, proliferation, antigen-presenting capacity and transcriptional reprogramming observed *in vitro*. However, by uncoupling temperature from the broader physiological component of fever, our localized heating model demonstrates that temperature alone is insufficient to enhance the humoral response following immunization. These findings establish intrinsic B cell thermosensitivity as a component of humoral immune regulation while indicating that its biological consequences depend on its integration within the multicellular and inflammatory signals that shape germinal center responses *in vivo*. More broadly, the experimental platform described here provides a unique opportunity to investigate the contribution of local temperature to immune responses independently of the multiple confounding factors that have previously limited such studies *in vivo*.

## Acknowledgments

The authors would like to thank Khadija Bertino and all the secretary staff of LVTS, and Thomas Lelant, Emma Sylvain-Plassard and Grégory Denjean from the LVTS mouse facility. AI (ChatGPT, Claude.ai) was used for text editing.

## Authors contributions

S.M. (Investigation [Lead], Methodology [Lead], Visualization [Lead], Writing—original draft [Lead], Writing—review & editing [Equal]), Z.S. (Investigation [Equal]), P.L. (Methodology [Equal], I.H. (Investigation [Equal]), S.H. (Investigation [Equal]), E.P. (Investigation [Equal]), O.M. (Investigation [Equal]), G.F. (Conceptualization [Equal], Writing—review & editing [Equal]), G.C. (Conceptualization [Equal], Writing—review & editing [Equal]), A.N. (Conceptualization [Equal], Writing—review & editing [Equal]), M.L.B. (Conceptualization [Lead], Funding acquisition [Lead], Investigation [Equal], Methodology [Equal], Supervision [Lead], Visualization [Equal], Writing—review & editing [Lead]).

## Abbreviations

FC: Flow cytometry
FRT: fever-range temperature
GC: germinal center
MFI: median fluorescence intensity
NP: 4-hydroxy-3-nitrophenylacetyl
SPAg: surrogate particulate antigen

