## Supplemental Figures and Tables for "Fever-range temperature enhances B cells *in vitro* but is dispensable for the germinal center *in vivo*"

**Running title:** Fever-range hyperthermia impact on B cell responses


#### Authors:

Sofiane Mohammed[1], Zineb Sbihi[1], Patrick Laignel[1], Ines Houtia[1], Sabrina Haddou[1], Emanuele Procopio[1], Océane Morales[1], Grégory Franck[1], Giusepina Caligiuri [1,2], Antonino Nicoletti[1], Marie Le Borgne[1*]

[1]Université Paris Cité and Université Sorbonne Paris Nord, INSERM, LVTS, F-75018 Paris, France

[2]Department of Cardiology, Assistance Publique-Hôpitaux de Paris, Hôpitaux Universitaires Paris Nord Val-de-Seine, Site Bichat, Paris, France

[*] Corresponding author

### Supplementary figures:

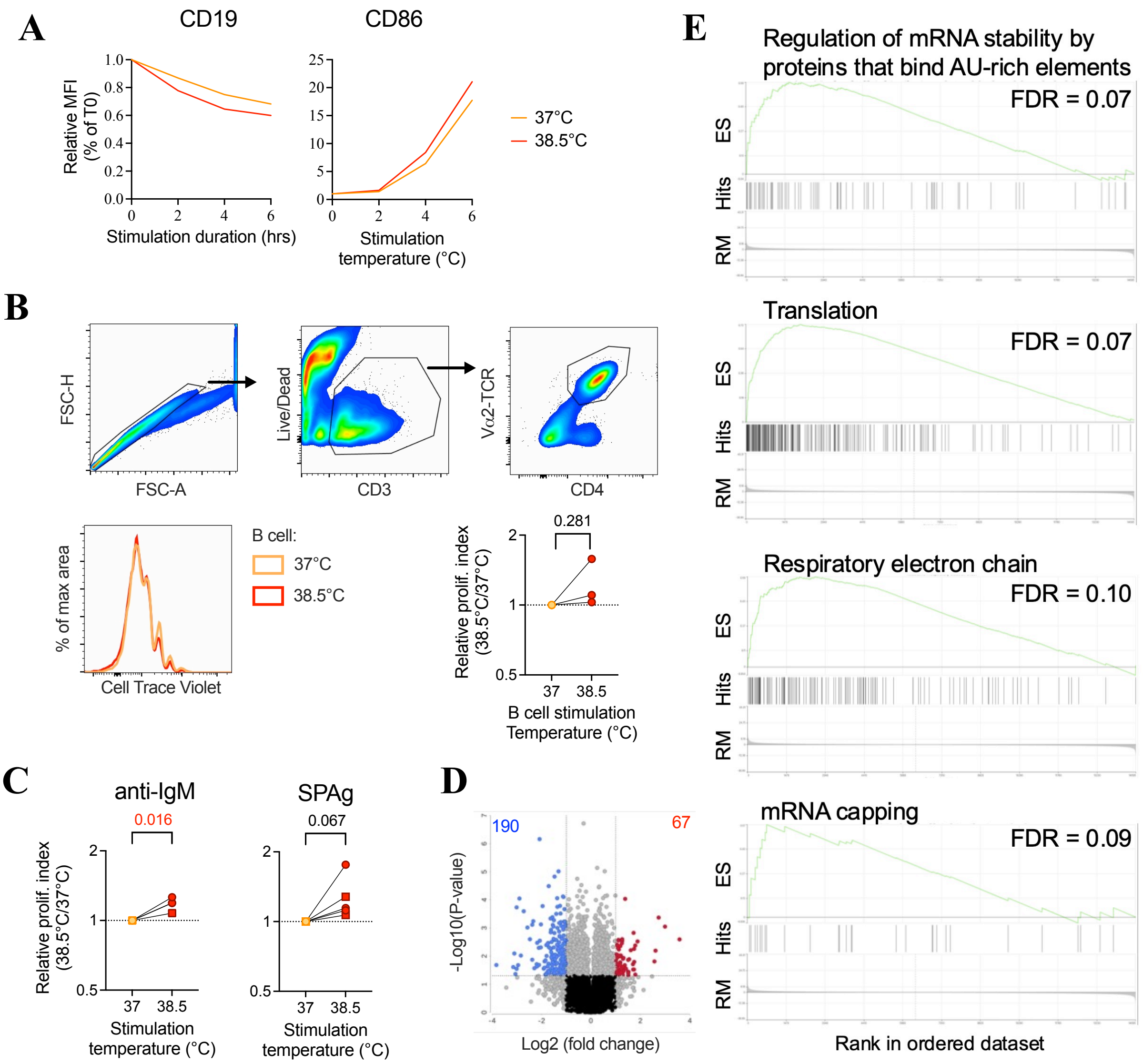


**Supplementary Figure 1. Characterization of B cell responses to moderate hyperthermia *in vitro*.** (A) Time course of surface CD19 and CD86 expression (MFI, median fluorescence intensity) on splenic B cells stimulated with SPAg-OVA + anti-CD40 at 37 °C versus 38.5 °C, measured at 0, 1, 2, 4, and 6 h by flow cytometry. Each point represents one independent experiment. Related to Figure 1A. (B) Representative gating strategy used to identify live, singlet CD4$^{+}$ OT-II T cells in the B cell–T cell co-culture assay, representative proliferation histogram, and corresponding T cell fold change in T cell proliferation index (mean number of divisions among divided cells, calculated from proliferation index – 1) at day 3. Each dot represents one independent experiment. Related to Figure 1B. (C) Proliferation index of splenic B cells following 24 h stimulation with either anti-IgM + anti-CD40 or SPAg-OVA + anti-CD40 at 37 °C or 38.5 °C. Data are presented as fold changes in proliferation index (calculated from proliferation index – 1) relative to the paired 37 °C condition. Each dot represents one independent experiment. Related to Figure 1C. Numbers indicate the p-values from one-sample t-tests performed on log2-transformed within-experiment ratios. P-values ≤ 0.05 are shown in red. (D) Volcano plot of differential gene expression (RNA-Seq) between B cells stimulated for 4 h with SPAg-OVA at 38.5 °C versus 37 °C. Significantly up- (red) and down-regulated (blue) genes (|log2FC| > 2, FDR <0.05) are highlighted. (E) Gene set enrichment plots for selected pathways significantly enriched at 38.5 °C versus 37 °C following GSEA analysis of RNA-seq data from B cells stimulated with SPAg-OVA for 4 h at 37 °C or 38.5 °C. False discovery rate (FDR) values are indicated for each pathway. GSEA was performed on Reactome and Gene Ontology pathways (FDR ≤ 0.1), using RNA-Seq data from n = 3 biological replicates per condition. ES: enrichment score; RM: ranking metric score.

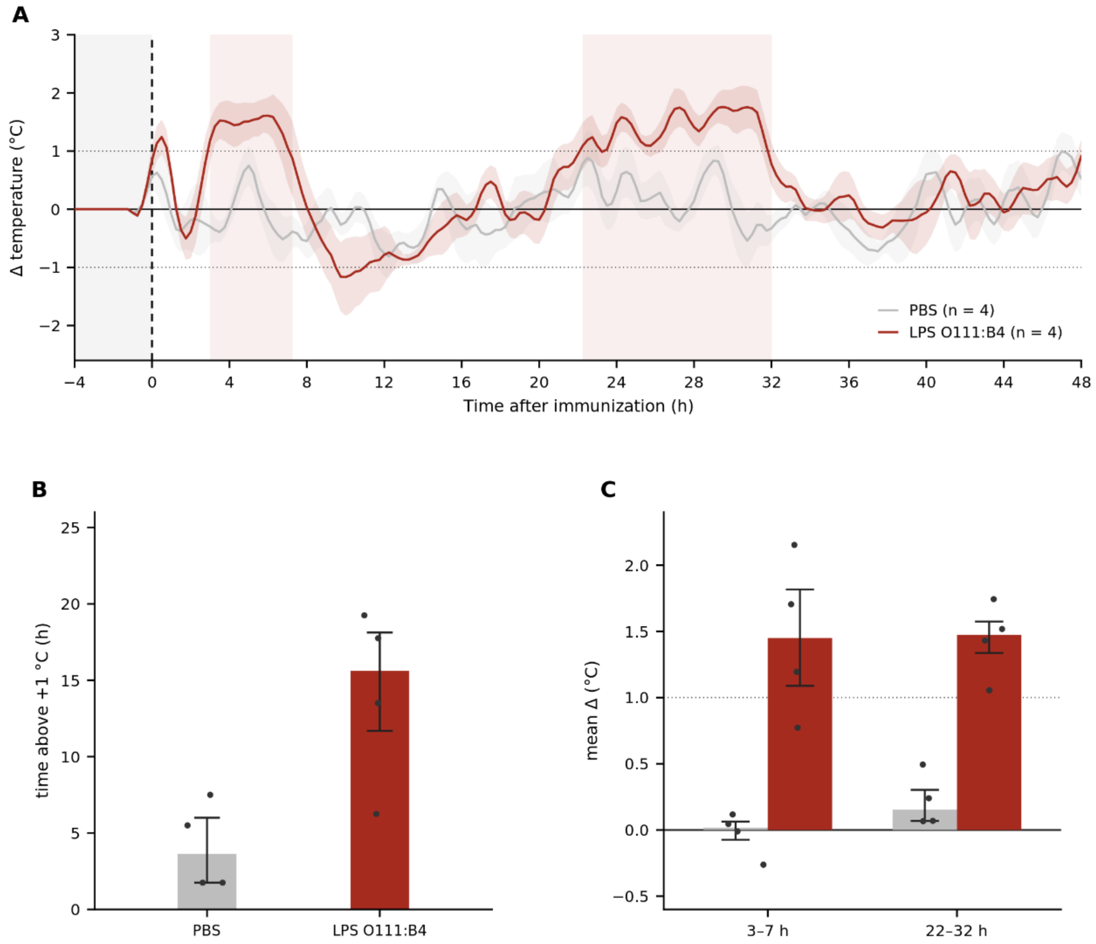


**Supplementary Figure 2. Subcutaneous LPS co-injected with NP-OVA induces a biphasic rise in core body temperature.** Core body temperature of wild-type C57BL/6 mice was recorded every 15 min by intraperitoneal RFID temperature microchips (UID). For each animal, temperature is expressed as the deviation from its own temperature at the same time point 24 h before the injection, and smoothed by a second-order Savitzky-Golay filter over a 2 h window. Mice received a subcutaneous hock injection of 50 µg LPS (~2 mg/kg) from Escherichia coli O111:B4 together with NP-OVA, or of PBS. (A) Group mean ± SEM from 4 h before to 48 h after injection; dotted lines mark ±1 °C and shaded bands indicate the two febrile phases defined in (C). (B) Time spent above +1 °C over the first 48 h, 15.6 h after LPS against 3.6 h after PBS. (C) The two febrile phases were defined as the intervals over which the group mean of the LPS-injected mice exceeds +1 °C, namely 3.0

to 7.3 h and 22.3 to 32.0 h. Mean temperature deviation of each mouse during each febrile phase reached +1.45 °C against +0.02 °C in the first phase and +1.47 °C against +0.15 °C in the second. Bars show the median, whiskers the interquartile range, and each point represents one mouse. n = 4 mice per group; Mann-Whitney test, $p = 0.029$ for both febrile phases and $p = 0.059$ for the time spent above +1 °C.

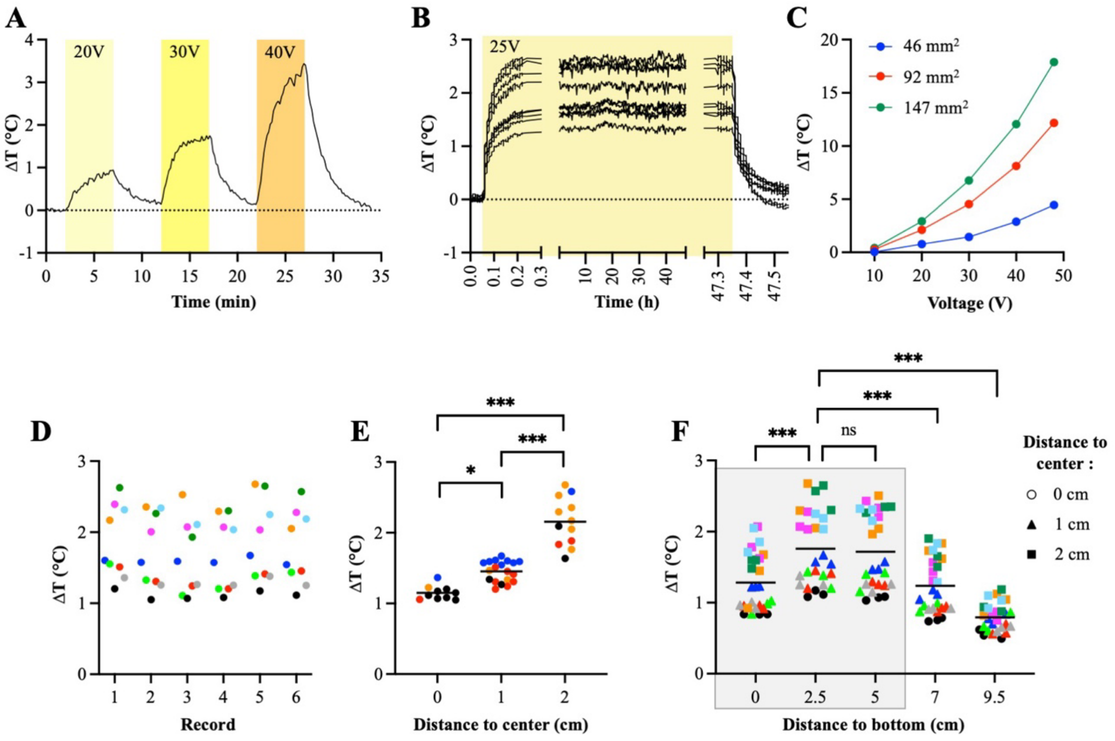


**Supplementary Figure 3.** *Ex vivo* characterization of the inductive heating properties of metallic objects. (A) Representative kinetics of metallic object temperature during repeated ON/OFF cycles of the electromagnetic field, measured ex vivo by infrared thermometry at increasing applied voltages (20, 30, 40V), as shown in video 1. (B) Stability of induced temperature over 48 h of continuous electromagnetic exposure (25 V). (C) Temperature increases as a function of metallic object size and applied voltage. Each point represents one object (one object per size), recorded simultaneously at identical positions. (D) Variability between acquisition movies, assessed from repeated recordings of multiple objects acquired under identical spatial configurations. (E) Effects of object identity and horizontal (x–y) distance from the center on heating, evaluated by exchanging object positions between recordings. (F) Effect of vertical (z) position on heating, assessed from recordings in which horizontal positions were fixed across objects. Temperatures in (A–B) are expressed as ΔT_washer ($t_x$–$t_0$) – ΔT_background ($t_x$–$t_0$). For (C–F), the maximum ΔT reached within

8 min was used. In (D–F), each point represents one object, color-coded by identity. Source of variations were analyzed using linear mixed-effects models fitted by REML, with distance (E) or vertical position (F) as fixed effects and experiments and objects as random effects; recording and object were treated as random effects for variance estimation in (D). Post-hoc comparisons were conducted when the global effect of position was significant

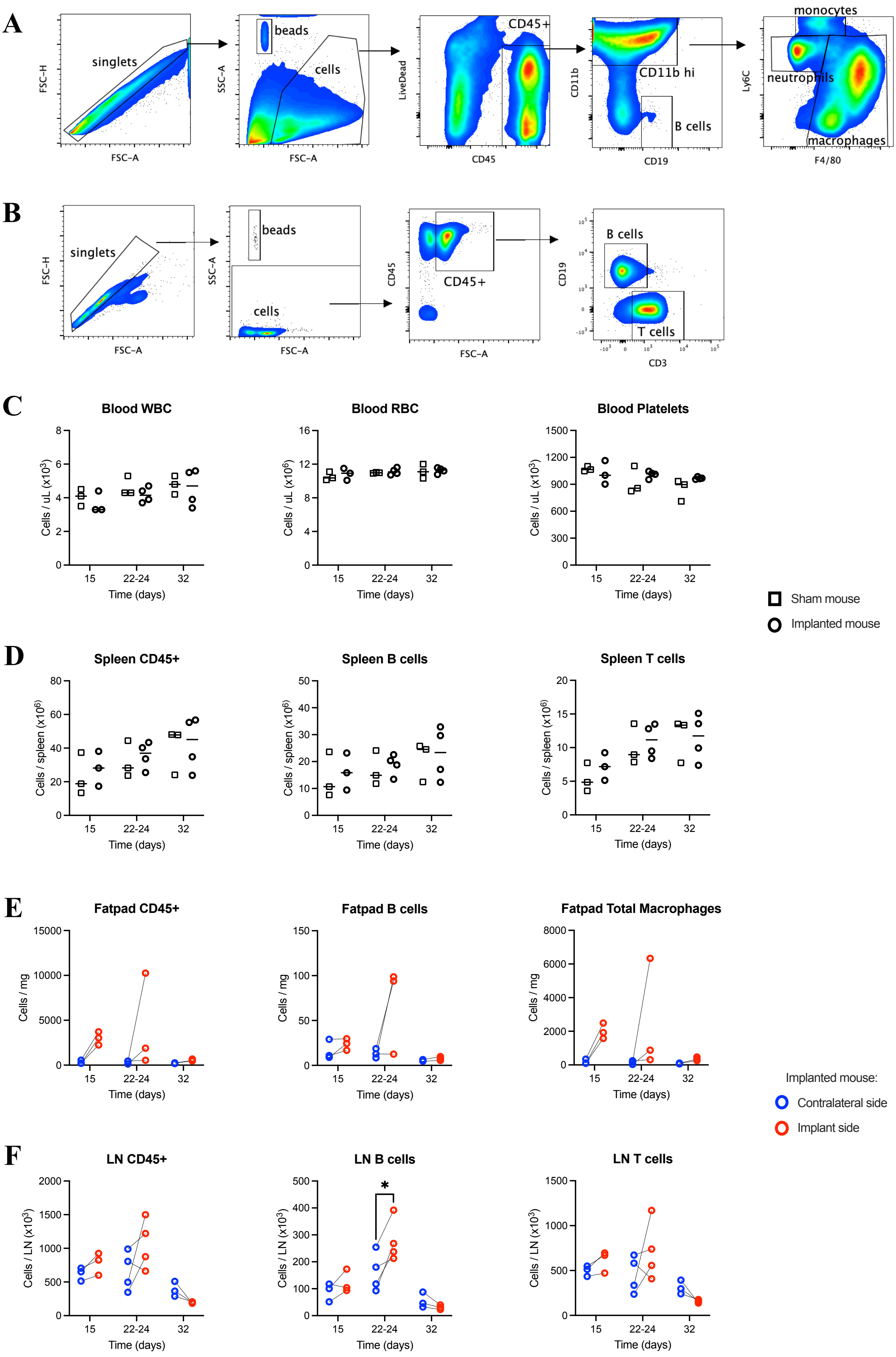
A
singlets
FSC-H
FSC-A
beads
cells
SSC-A
LiveDead
CD45
CD45+
CD11b
CD11b hi
CD19
B cells
Ly6C
monocytes
neutrophils
macrophages
F4/80
B
singlets
beads
cells
CD45+
CD3
T cells
C
Blood WBC
Blood RBC
Blood Platelets
Cells / uL (x10$^3$)
Cells / uL (x10$^6$)
Time (days)
15
22-24
32
Sham mouse
Implanted mouse
D
Spleen CD45+
Spleen B cells
Spleen T cells
Cells / spleen (x10$^6$)
E
Fatpad CD45+
Fatpad B cells
Fatpad Total Macrophages
Cells / mg
Implanted mouse:
Contralateral side
Implant side
F
LN CD45+
LN B cells
LN T cells
Cells / LN (x10$^3$)
*

**Supplementary Figure 4. The metallic implant does not disrupt systemic or local immune homeostasis.** Mice were implanted with a metallic object around one inguinal lymph node. Leucocyte populations in the blood, spleen, lymph node and fatpad around the inguinal lymph node (contralateral side and implant side) were analyzed by flow cytometry after 15 to 32 days. (A) Gating strategy in the fatpad. (B) Gating strategy in the spleen and LN. (C) White blood cell (WBC), red blood cell (RBC), and platelet counts in blood over time post-implantation in sham and implanted mice. (D) Absolute counts of $CD45^+$, B ($CD19^+$) and T ($CD3^+$) cells in the spleen over time post-implantation. (E) Absolute counts of $CD45^+$ cells, B cells and macrophages in the perilymphatic inguinal adipose tissue of the implant and contralateral sides over time in implanted mice. (F) Absolute counts of $CD45^+$, B and T cells in the inguinal lymph node of the implant and contralateral sides over time in implanted mice.. A transient increase observed at approximately 3 weeks post-implantation is fully resolved by week 4. Each point represents one mouse. *: $p<0.01$, mixed-effects model followed by Sidak's test.

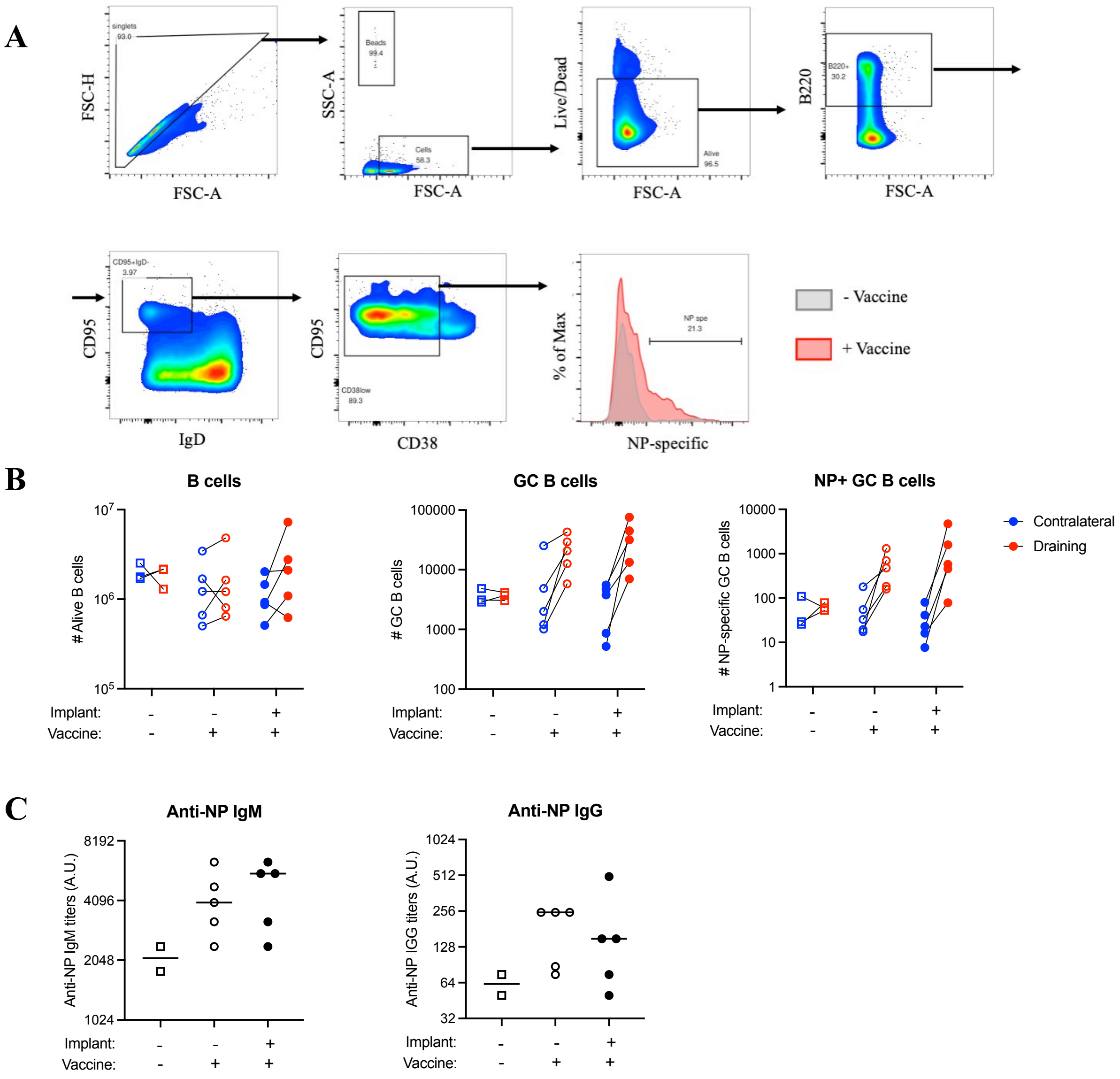


**Supplementary Figure 5. The metallic implant does not impair lymphatic drainage or the establishment of a primary humoral response during active immunization.** Wild-type female C57BL/6N mice (n = 13) were assigned to three groups: sham-operated and injected with PBS (ctrl, n = 3), sham-operated and immunized (n = 5), or implanted with a metallic ring around the left inguinal lymph node and immunized (n = 5). At D0, mice received a subcutaneous hock injection of 20 µL containing either PBS alone or 50 µg NP-OVA together with 1 µg LPS from *Escherichia coli* O111:B4. LPS dose 1 µg (40 to 50 µg/kg) does not elicit a febrile response, so that drainage and priming could be assessed independently of any thermal effect. No electromagnetic field was applied. Draining and contralateral inguinal lymph nodes and serum were collected at D9. (A) Gating strategy. (B) Absolute number of live cells (left), of germinal center B cells (B220$^{+}$ CD19$^{+}$ IgD$^{-}$ CD95$^{+}$ CD38$^{low}$, middle) and of NP-specific germinal center B cells (same gate, NP$^{+}$, right) in the draining (red) and contralateral (blue) inguinal lymph node, on a logarithmic scale. Lines connect the two dots of the same mouse. (C) Serum anti-NP IgM (left) and anti-NP IgG (right) titers at D9. Each dot represents one mouse; in (C) horizontal bars indicate the median.

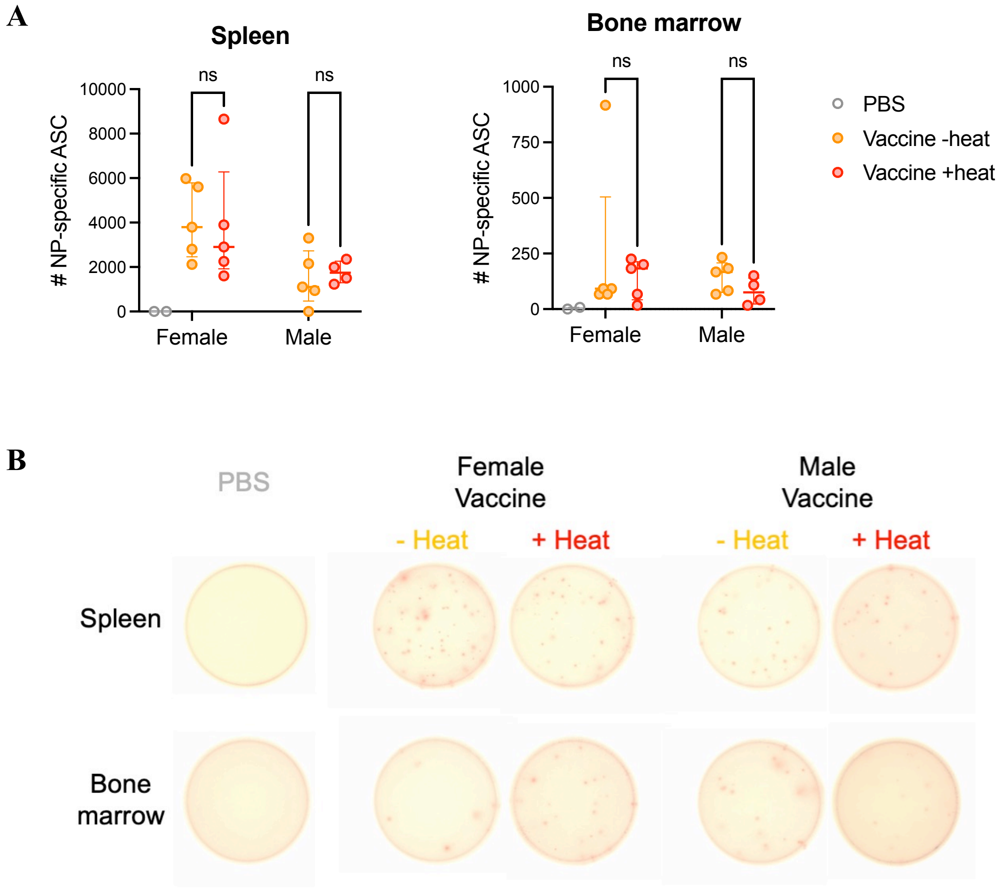


**Supplementary Figure 6. Effect of initial localized hyperthermia on antibody-secreting cells after a challenge.** (A) Number of NP-specific IgG-secreting antibody-secreting cells (ASCs) per spleen (left) and per tibia bone marrow (right) at D28, seven days after the boost, measured by ELISPOT on serial fivefold dilutions in a subset of mice from the recall cohort (batch 2 of Figure 4). Females and males are shown separately. PBS (grey), Vaccine − Heat (orange), Vaccine + Heat (red). (B) Representative ELISPOT wells for each condition, spleen (top row) and bone marrow (bottom row). Each point represents one mouse and bars indicate the median with the interquartile range. NP-specific ASCs were undetectable in PBS-injected mice in both organs, confirming the specificity of the assay. Immunized groups were compared by two-way ANOVA (sex × heat), followed by Šídák's multiple comparison test between heated and non-heated mice within each sex (n = 4 to 5 mice per group and sex); PBS-injected mice were not included in the analysis. Early localized hyperthermia had no effect on the number of NP-specific ASCs in either organ, in females or in males (ns).

#### Supplementary Movies:

**Movie 1. Heating of ring-shaped metallic objects *ex vivo*.** Four metallic objects (indicated by arrows) were placed in the heating system (distance to bottom: 2,5 cm). Repeated ON/OFF cycles of the electromagnetic field were performed and measured by infrared thermometry at increasing applied voltages (20, 30, 40V). Time is indicated in min:s ; the temperature scale is shown on the right of the video.

**Movie 2. Heating of ring-shaped metallic objects in mice.** Ring-shaped metallic objects were implanted s.c. in the back of two mice. Three weeks later, the mice were placed in the heating system, and infrared thermal imaging was performed. After 2 min 30 s, the electromagnetic field was turned on (40 V) for 7 min 30 s and then turned off, allowing the objects to cool down. Time is indicated in min:s ; the temperature scale is shown on the right of the video.

### Supplementary Tables:

**Supplementary Table 1.** Reagents, consumables and kits.

| Product | Supplier | Catalogue number | Working concentration |
|---|---|---|---|
| **In vivo experiments** | | | |
| Alhydrogel adjuvant 2% | InvivoGen | vac-alu-50 | — |
| LPS from Escherichia coli O111:B4 | Sigma | L4391 | — |
| NP-OVA | LGC BioSearch | N-5051 | — |
| Temperature Programmable Microchip 2.12 × 11.5 | UID Identification Solutions | UCT-2112 | — |
| **Flow cytometry and tissue preparation** | | | |
| Cell Strainer 100 µm | Corning | 431752 | — |
| Cell Strainer 70 µm | Corning | 431751 | — |
| PBS 10X (for tissue preparation) | Gibco | 14200-083 | — |
| PBS 1X | Gibco | 14190-169 | — |
| EDTA 0.5M | Invitrogen | 15575-038 | — |
| MACS BSA Stock Solution | Miltenyi Biotec | 130-091-376 | — |
| IMDM with Glutamax | Gibco | 31980-048 | — |
| gentleMACS C tubes | Miltenyi Biotec | 130-093-237 | — |
| Collagenase D from Clostridium histolyticum | Roche | 11088858001 | 0.1 U/mL |
| DNase I grade II, from bovine pancreas | Roche | 10104159001 | 20 U/mL |
| Brilliant Stain Buffer | BD Biosciences | 566349 | 1/4 |
| Purified Rat Anti-Mouse CD16/CD32 (Mouse BD Fc Block) | BD Biosciences | 553142 | 1/500 |
| Precision Count Beads | Biolegend | 424902 | — |
| **ELISA/ELISPOT** | | | |
| high-binding 96-well plates (ELISA) | Greiner | 655061 | — |
| MultiScreen-IP plates (ELISPOT) | Millipore | MSIPS4W10 | — |

| Product | Supplier | Catalogue number | Working concentration |
|---|---|---|---|
| PBS 1X | Gibco | 14190-169 | — |
| Ultrapure BSA | Sigma-Aldrich | A3803 | — |
| Tween 20 | Sigma-Aldrich | P7949 | — |
| NP2-BSA | LGC Biosearch | N-5050XL | — |
| NP14-BSA | LGC Biosearch | N-5050M | — |
| Goat anti-Mouse IgM-HRP | Invitrogen | PA1-84383 | 1/2000 |
| Goat Anti-Mouse IgG-HRP | Southern Biotech | 1032-05 | 1/4000 |
| Donkey anti-Mouse IgG HRP | Invitrogen | SA1-100 | 1/2000 |
| TMB Chromogen | Thermo Scientific | N301 | — |
| ELISA Stop Solution | Thermo Scientific | N600 | — |
| AEC Chromogen/Substrate | BD Biosciences | 551951 | — |
| **Cell culture** | | | |
| RPMI 1640 + Glutamax | Gibco | 61870-044 | — |
| HEPES 1M | Gibco | 15360-130 | — |
| sodium pyruvate 100 mM | Gibco | 11360-039 | — |
| penicillin/streptomycin | Gibco | 15070-063 | — |
| 2-mercaptoethanol (50 mM) | Gibco | 31350-010 | — |
| CD19 MicroBeads, mouse | Miltenyi Biotec | 130-121-301 | — |
| LS columns | Miltenyi Biotec | 130-042-401 | — |
| Dynabeads™ Mouse CD43 (Untouched™ B Cells) kit | Invitrogen | 11422D | — |
| Dynabeads™ Untouched Mouse CD4 Cells Kit | Invitrogen | 11415D | — |
| MACS BSA Stock Solution | Miltenyi Biotec | 130-091-376 | — |
| mouse IL-4 | R&DSystems | 404-ML | — |
| mouse IL-5 | Biolegend | 581506 | — |
| Rat anti-Mouse CD40, clone 1C10, functional grade | eBiosciences | 16-0401-85 | — |
| F(ab')2-Goat anti-Mouse IgM, Functional Grade | eBiosciences | 16-5092-85 | — |
| CellTrace Violet proliferation kit | Invitrogen | C34557 | — |
| **SPAg preparation** | | | |

| Product | Supplier | Catalogue number | Working concentration |
|---|---|---|---|
| Zeba™Spin Desalting Columns 7K MWCO | Thermo Scientific | 89894 | — |
| EZ-Link™ Sulfo NHS-LC-LC-Biotin | Thermo Scientific | A35358 | — |
| Purified Rat Anti-Mouse Ig, κ Light Chain, clone 187.1 | BD Biosciences | 559749 | — |
| PBS 1X | Gibco | 14190-169 | — |
| Ovalbumin | Sigma-Aldrich | A5503 | — |
| Bovine Serum Albumin (BSA) | Sigma-Aldrich | A7906 | — |
| Ultrafree® Centrifugal Filter 0.1 µm, PVDF | Millipore | UFC30VV00 | — |
| Ultrafree® Centrifugal Filter 0.65 µm, PVDF | Millipore | UFC30DV00 | — |
| MACS BSA Stock Solution | Miltenyi Biotec | 130-091-376 | — |
| streptavidin coated microspheres (660, 690), 0.4 µm | Bangs laboratories | CP01F | — |
| Anti-Ovalbumin antibody (FITC) | Abcam | ab85584 | 1/100 |
| Goat anti-mouse IgG-BV421, polyclonal | Biolegend | 405317 | 1/100 |
| **Transcriptomics** | | | |
| RNeasy Micro Kit | Qiagen | 74004 | — |
| TruSeq Stranded mRNA Library Prep | Illumina | 20020595 | — |

**Supplementary Table 2.** Antibody panels used for flow cytometry.

| Target | Fluorochrome | Clone | Supplier | Catalogue number | Dilution |
|---|---|---|---|---|---|
| **OT-II T cell purity** | | | | | |
| Vα2 TCR | BUV395 | B20.1 | BD Biosciences | 743834 | 1/200 |
| CD3 | BV421 | 17A2 | BD Biosciences | 564008 | 1/200 |
| Vβ 5.1, 5.2 TCR | BV650 | NR9-4 | BD Biosciences | 743001 | 1/200 |
| CD4 | PE-Cy7 | RM4-5 | BD Biosciences | 552775 | 1/400 |
| Viability (LIVE/DEAD Fixable) | Near-IR | — | Thermo Fisher | L34976 | 1/1000 |
| **B cell purity** | | | | | |
| CD 62 L | BUV395 | MEL-14 | BD Biosciences | 740218 | 1/200 |
| CD43 | BV421 | S7 | BD Biosciences | 562958 | 1/200 |
| CD19 | Super Bright 600 | 1D3 | eBioscience | 63-0193-82 | 1/200 |
| CD23 | BV711 | B3B4 | BD Biosciences | 563987 | 1/200 |
| IgM | PE | R6.60-2 | BD Biosciences | 553409 | 1/200 |
| CD5 | APC | 54-7.3 | BD Biosciences | 550035 | 1/200 |
| Viability (LIVE/DEAD Fixable) | Near-IR | — | Thermo Fisher | L34976 | 1/1000 |
| **In vitro B cell activation** | | | | | |
| CD 62 L | BUV395 | MEL-14 | BD Biosciences | 740218 | 1/200 |
| CD19 | BUV737 | 1D3 | BD Biosciences | 612781 | 1/200 |
| CD86 | BV421 | GL1 | BD Biosciences | 564198 | 1/200 |
| Ig, λ1, λ2 & λ3 Light Chain | BV650 | R26-46 | BD Biosciences | 744526 | 1/200 |
| I-A$^b$ MHC | FITC | AF6-120.1 | BD Biosciences | 553551 | 1/200 |
| CD95 | PE-Cy7 | Jo2 | BD Biosciences | 557653 | 1/200 |
| SPAg (anti-Igκ 187.1 on streptavidin microspheres) | APC | — | Bangs laboratories | CP01F | — |

| Target | Fluorochrome | Clone | Supplier | Catalogue number | Dilution |
|---|---|---|---|---|---|
| Viability (LIVE/DEAD Fixable) | Near-IR | — | Thermo Fisher | L34976 | 1/1000 |
| **In vitro B cell proliferation** | | | | | |
| CD38 | BUV395 | 90/ CD38 | BD Biosciences | 740245 | 1/200 |
| CD19 | BUV737 | 1D3 | BD Biosciences | 612781 | 1/200 |
| Proliferation tracking dye | CellTrace Violet | — | Invitrogen | C34557 | — |
| IgD | SB600 | 11-26C | eBioscience | 63-5993-82 | 1/400 |
| Ig, λ1, λ2 & λ3 Light Chain | BV650 | R26-46 | BD Biosciences | 744526 | 1/200 |
| GL7 | PE | GL7 | BioLegend | 144608 | 1/200 |
| CD95 | PE-Cy7 | Jo2 | BD Biosciences | 557653 | 1/200 |
| SPAg (anti-Igκ 187.1 on streptavidin microspheres) | APC | — | Bangs laboratories | CP01F | — |
| Viability (LIVE/DEAD Fixable) | Near-IR | — | Thermo Fisher | L34976 | 1/1000 |
| **Antigen presentation, OT-II T cell proliferation** | | | | | |
| Proliferation tracking dye | CellTrace Violet | — | Invitrogen | C34557 | — |
| CD19 | BV605 | 6D5 | BioLegend | 115539 | 1/200 |
| CD25 | BV711 | PC61 | BD Biosciences | 740714 | 1/200 |
| CD3 | FITC | 145-2C11 | BD Biosciences | 553062 | 1/200 |
| CD69 | PE | H1.2F3 | BD Biosciences | 553237 | 1/200 |
| Vα2 TCR | PerCP-Cy5.5 | B20.1 | BD Biosciences | 560529 | 1/200 |
| CD4 | PE-Cy7 | RM4-5 | BD Biosciences | 552775 | 1/400 |
| SPAg (anti-Igκ 187.1 on streptavidin microspheres) | APC | — | Bangs laboratories | CP01F | — |
| Viability (LIVE/DEAD Fixable) | Near-IR | — | Thermo Fisher | L34976 | 1/1000 |

| Target | Fluorochrome | Clone | Supplier | Catalogue number | Dilution |
|---|---|---|---|---|---|
| **In vivo B cell response** | | | | | |
| CD19 | BUV737 | 1D3 | BD Biosciences | 612781 | 1/200 |
| IgM | BV421 | R6-60.2 | BD Biosciences | 562595 | 1/200 |
| IgD | SB600 | 11-26C | eBioscience | 63-5993-82 | 1/400 |
| CD138 | BUV395 | 281-2 | BD Biosciences | 740240 | 1/200 |
| IgG1 | APC | X56 | BD Biosciences | 550874 | 1/200 |
| CD95 | PE-Cy7 | Jo2 | BD Biosciences | 557653 | 1/200 |
| CD38 | BUV805 | 90/ CD38 | BD Biosciences | 741955 | 1/200 |
| NP-BCR | PE | chemical conjugate | LGC BioSearch | N-5070 | 1/200 |
| Viability (LIVE/DEAD Fixable) | Near-IR | — | Thermo Fisher | L34976 | 1/1000 |
| **In vivo main leukocyte subsets** | | | | | |
| CD19 | BUV737 | 1D3 | BD Biosciences | 612781 | 1/500 |
| CD3 | eFluor450 | 17A2 | eBioscience | 48-0032-82 | 1/200 |
| CD45 | BV605 | 30- F11 | BD Biosciences | 563053 | 1/200 |
| CD 11b | BV711 | M1-70 | BioLegend | 101241 | 1/200 |
| CD4 | FITC | GK1.5 | BD Biosciences | 553729 | 1/200 |
| CD193 | PE-Cy7 | J073E5 | BioLegend | 144514 | 1/200 |
| Ly6C | APC | HK1.4 | BioLegend | 128016 | 1/200 |
| Ly6G | A700 | 1A8 | BD Biosciences | 561236 | 1/200 |
| CD8 | APC-Cy7 | 53-6.7 | BD Biosciences | 557654 | 1/200 |
| **In vivo fat pad analysis** | | | | | |
| CD19 | BUV737 | 1D3 | BD Biosciences | 612781 | 1/500 |
| I-A/I-E | BV421 | M5/114.15.2 | BioLegend | 107631 | 1/500 |
| CD45 | BV605 | 30- F11 | BD Biosciences | 563053 | 1/200 |
| CD 11b | BV711 | M1-70 | BioLegend | 101241 | 1/200 |
| F4/80 | A488 | BM8 | BioLegend | 123120 | 1/200 |

| Target | Fluorochrome | Clone | Supplier | Catalogue number | Dilution |
|---|---|---|---|---|---|
| CD3 | PE | 145-2C11 | BD Biosciences | 553064 | 1/100 |
| Ly6C | APC | HK1.4 | BioLegend | 128016 | 1/200 |
| Ly6G | A700 | 1A8 | BD Biosciences | 561236 | 1/200 |
| Viability (LIVE/DEAD Fixable) | Near-IR | — | Thermo Fisher | L34976 | 1/1000 |
